# A novel image-based approach for soybean seed phenotyping using machine learning techniques

**DOI:** 10.1101/2022.10.10.511645

**Authors:** Melissa Cristina de Carvalho Miranda, Alexandre Hild Aono, José Baldin Pinheiro

## Abstract

Soybean is one of the most important sources of protein and vegetable oil in the world. Given its increasing demand, the increment in yield has only been possible due to investments in research and production technology, especially in breeding programs. One of the main factors influencing soybean yield is the seed morphology; however, its analyses are hampered by the lack of efficient computational approaches with not only accurate results, but also a high flexibility to user preferences and needs. In this context, the present work provides a methodological framework for: (i) seed segmentation in soybean images; (ii) seed morphological evaluation; and (iii) image-based prediction of the hundred-seed weight trait. We used genotypes from a partial diallel cross design, which aimed at obtaining genotypes with high agronomic performance. In addition to the measurement of the hundred-seed weight, we collected RGB images of seeds of each plot. For image segmentation, we created an in-house image processing pipeline, which enabled a full morphological seed evaluation. For predicting the hundred-seed weight, we compared different machine learning algorithms using as input the morphological characteristics obtained, and also features from state-of-the-art convolutional neural network (CNN) architectures. The image segmentation methodology showed to be highly efficient, as more than 98% of the seeds in the images were correctly identified. Even if the seeds were close, the segmentation strategy could separate them into independent image components. In addition to supplying a highly accurate decision support system for soybean breeders, we verified the morphological phenotyping adaptability in other plant species, fully assessing the pipeline generalization. We consider the use of this methodology highly advantageous, as the method is entirely based on widely used morphological operations, which results in an easy implementation and low computational costs. Using these morphological measures, we could estimate machine learning models for predicting the hundred-seed weight, achieving considerable predictive accuracy. The same results were observed for CNN-obtained features, showing the efficiency of the morphological measurements as feature extractors. The possibility of obtaining seed morphological characteristics provides a valuable tool for the continuous and efficient development of new soybean cultivars in breeding programs aimed at long-term genetic gain. Additionally, through a faster seed image acquisition workflow, with less chance of errors and low cost, it is also possible to make predictions of important soybean characteristics. The work conducted has the potential to help future research and the industry to develop automated phenotyping tools, incorporating the proposed analytical workflows.

## 1 Introduction

As the most important source of protein and vegetable oil in the world, soybean (*Glycine max* L.) plays a significant role in global food security (Hartman et al., 2011). Soybean yield is a complex quantitative trait influenced by several factors, including the number of plants per unit area, the number of seeds per plant, and the weight of one hundred seeds (HSW), which is directly dependent on the size and shape of seeds (length, width and thickness) (Niu et al., 2013). For this reason, seed conformation is closely related to soybean yield and quality, being an important target in plant breeding according to the current demand (Panthee et al., 2005; Niu et al., 2013).

Seed morphological phenotypes are associated with several important soybean characteristics, including growth and development, physiology, biochemistry, and genetics (Baek et al., 2020). Although one of the main drivers of soybean yield expansion in recent decades has been the elevation in the number of seeds per plant, there have been no changes in the size of individual seeds over time (Ainsworth et al., 2012). Hu et al. (2013) point out that increasing seed size may be the next component explored to increase yield when the number of seeds per plant is fixed. Thus, the development of tools for the study of seed morphology represents a valuable resource in breeding programs, especially in crops for which the seed is the main commercial product, such as soybean.

Seed evaluation through visual inspections is one of the oldest techniques of phenotyping (Gustin and Settles, 2016), and archaeological evidence suggests that seed size was one of the first characteristics to suffer selective pressure in early crops (Purugganan and Fuller, 2009). Manual measurements using calipers have been the most adopted method for this task over the last few years, but it is a laborious, time-consuming, and extremely error-prone process (Li et al., 2019; Liu et al., 2020; Yang et al., 2021; Marrano and Moyers, 2022). This challenge of testing large and complex genetic populations under different environmental conditions prevented seed morphology from being accurately incorporated into breeding programs (Baek et al., 2020; Marrano and Moyers, 2022).

In this context, it is noteworthy that there is still an enormous need to find tools to obtain high quality seed phenotypic data, with fast, accurate, and non-destructive approaches, also including acceptable costs and repeatedly in entire populations (Liu et al., 2020). Thus, high-throughput phenotyping (HTP) emerged, successfully integrating plant science, engineering and computing (Shakoor et al., 2017). Through images of seeds obtained by different optical sensors, it is possible to measure their size and conformation, evaluating the plant response to biotic and abiotic stresses, as well as measuring their chemical composition (Gustin and Settles, 2016; Rahman et al., 2016; Momin et al., 2017).

The main application of HTP is to use secondary characteristics related to grain yield, resistance to biotic and abiotic stresses, and final quality that can be useful in performance tests of lines in breeding programs (Crossa et al., 2017). Indirect selection may be preferable when the secondary characteristic is less expensive to measure than the primary characteristic, accelerating the decision-making stages (Bernardo, 2010). Thus, it is assumed that through seed images it is possible to make predictions of important characteristics of the crop, such as the production component HSW. However, limited flexibility and poor performance of classical image processing pipelines for complex phenotyping tasks have been observed, and as a consequence machine learning (ML) techniques are expected to play a prominent role in the future of image-based phenotyping (Ubbens and Stavness, 2017; Koh et al., 2021; Van Dijk et al., 2021).

Convolutional neural networks (CNNs) are a class of deep learning methods that have become the state-of-the-art in many computer vision tasks (Van Dijk et al., 2021). Most computer vision applications address the red-green-blue visible spectrum (RGB, 400 – 700 nm) and should be able to identify and classify seeds based on external characteristics such as size, shape, color, and texture (Rahman et al., 2016). In contrast to classic image analysis approaches, which first measure the statistical properties of the image as features to be used to learn a data model, CNNs actively learn a variety of filter parameters during model training (Ubbens and Stavness, 2017). The true power of deep learning, especially for image recognition, comes from convolutional layers, which use the convolution and activation operation to efficiently extract features from the input data (Traore et al., 2018).

In the present study, we propose a soybean image segmentation method, combining mathematical morphological operations with traditional image segmentation methods. In addition to presenting high accuracy in seed segmentation in a wide range of images, we verified the adaptability of the method to carry out automatic segmentation of other plant species, in order to assess the generalizability of the methodology for different seed morphologies. Additionally, we investigate the potential of traditional ML models and CNNs to predict the HSW in soybean images, in order to improve the decision support system of soybean breeders.

## 2 Material and Methods

The present study was composed of three main steps (Fig. 1): (1) soybean population definition and image acquisition; (2) seed segmentation and characterization; and (3) image-based prediction via ML. In step (1), we obtained seed images for genotypes planted in contrasting environments for soybean Asian rust (Fig. 1-1a). These plants were evaluated for the HSW trait, and in addition we captured RGB pictures for every genotype using 100 seeds in each image considering different seed distributions on the scanner (Fig. 1-1d). In (2), we propose an automated seed segmentation methodology through the combination of mathematical morphological operations and Otsu’s thresholding, supplying a robust approach for soybean phenotyping with high accuracies in images with densely distributed seeds (Fig. 2-2). Based on the segmentation process, we calculated 6 different seed measurements, also evaluating their potential as a complementary method for phenotypic evaluation (Fig. 2-2j). In addition to soybean seeds, we also tested the feasibility of the approach developed for other crops (common bean, maize, and chickpea). In (3), we contrasted several ML-based algorithms for predicting HSW based on: (a) morphological characteristics obtained in (2) (Fig. 2–3a); and (b) image features obtained through different architectures of CNNs (Fig. 2–3b).

**Figure 1.**
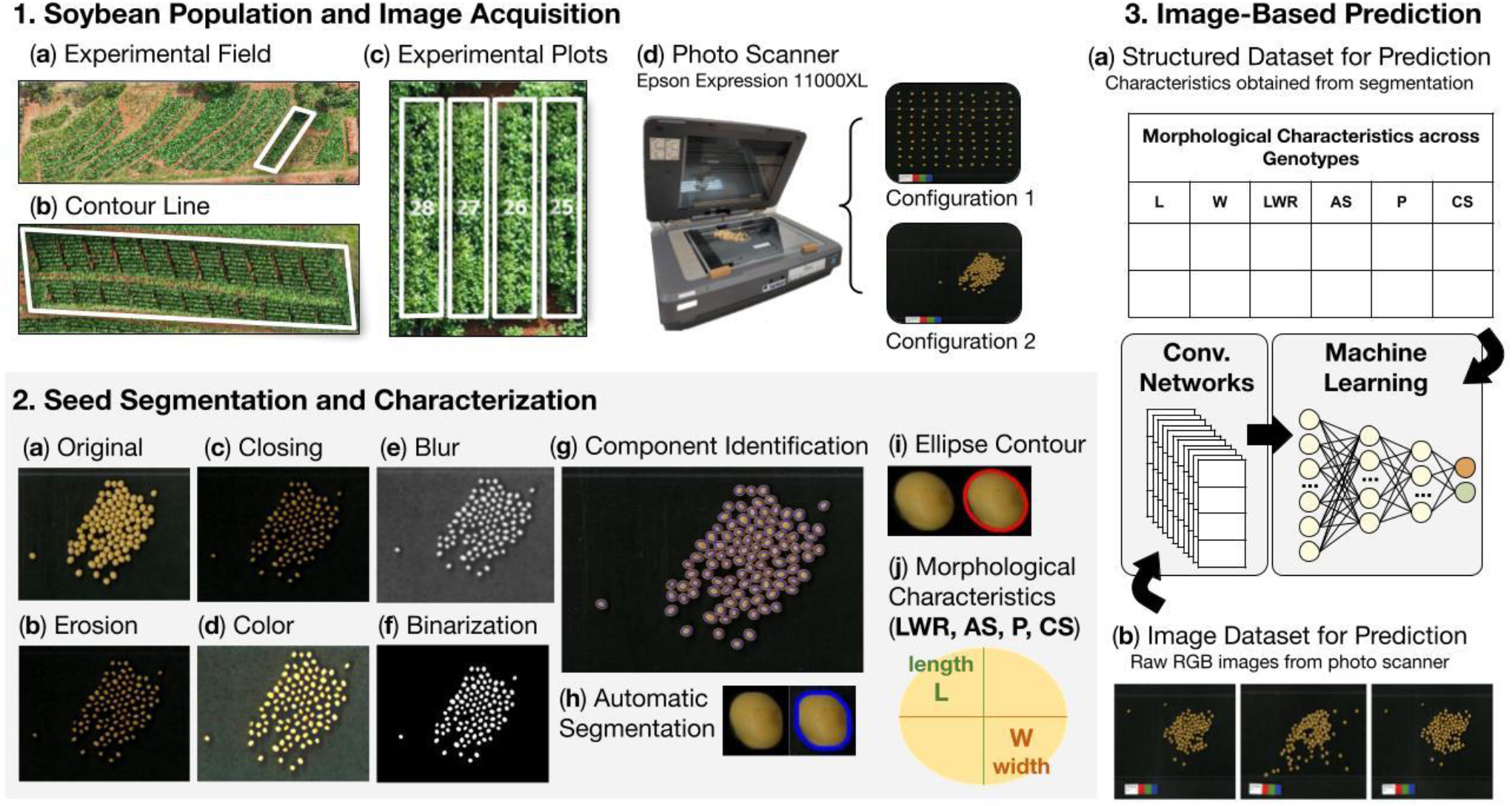
Overview of the process of obtaining seeds from the population of soybean genotypes (1a - 1c), acquisition of seed images (1d), image segmentation (2a - 2i), calculation of seed morphological characteristics (2j) and predictions based on ML models (3a – 3b).

**Figure 2.**
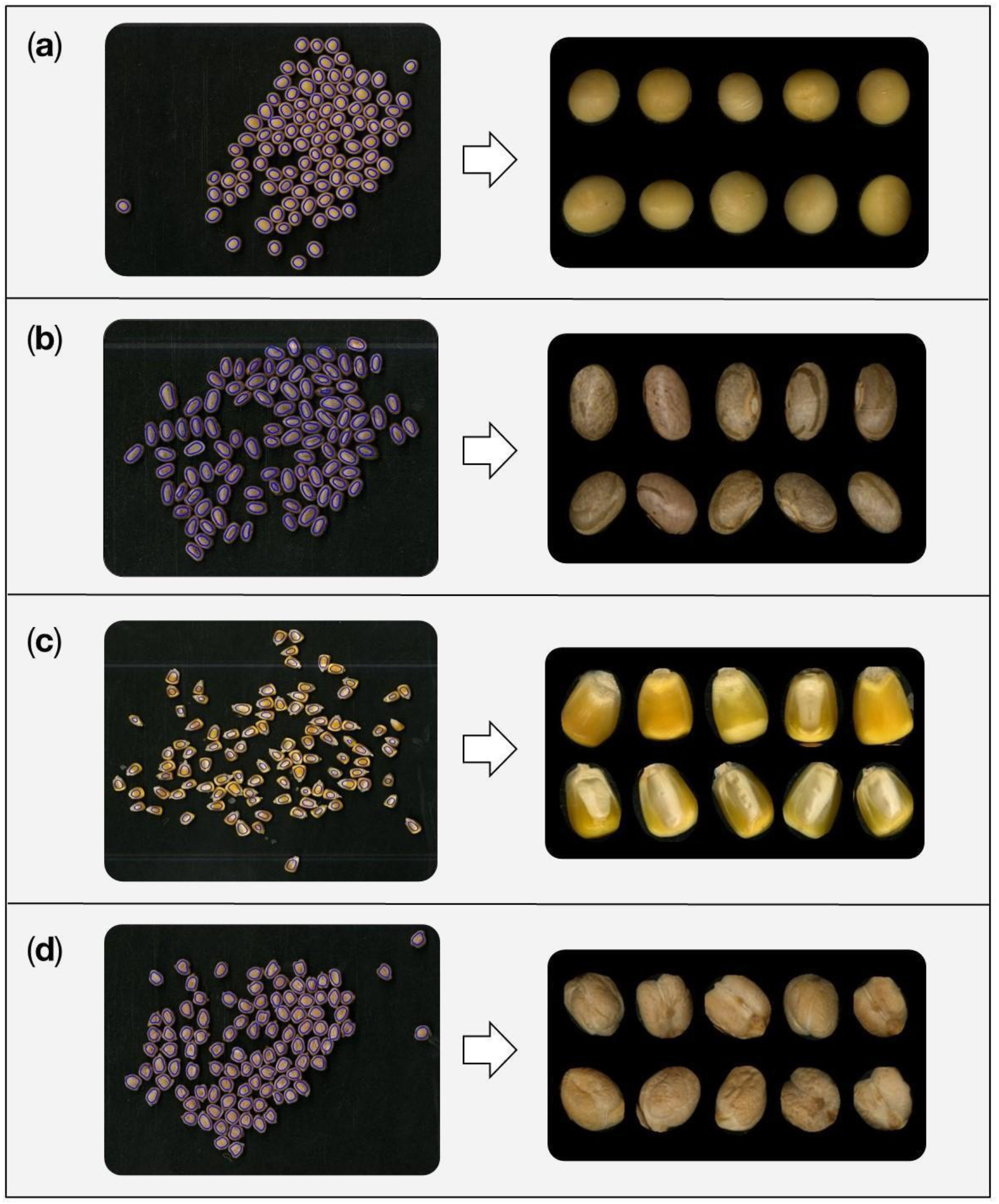
Identification of components in the images and segmentation result for seeds of (a) *Glycine max*, (b) *Phaseolus vulgaris*, (c) *Zea mays*, and (d) *Cicer arietinum*.

**Figure 3.**
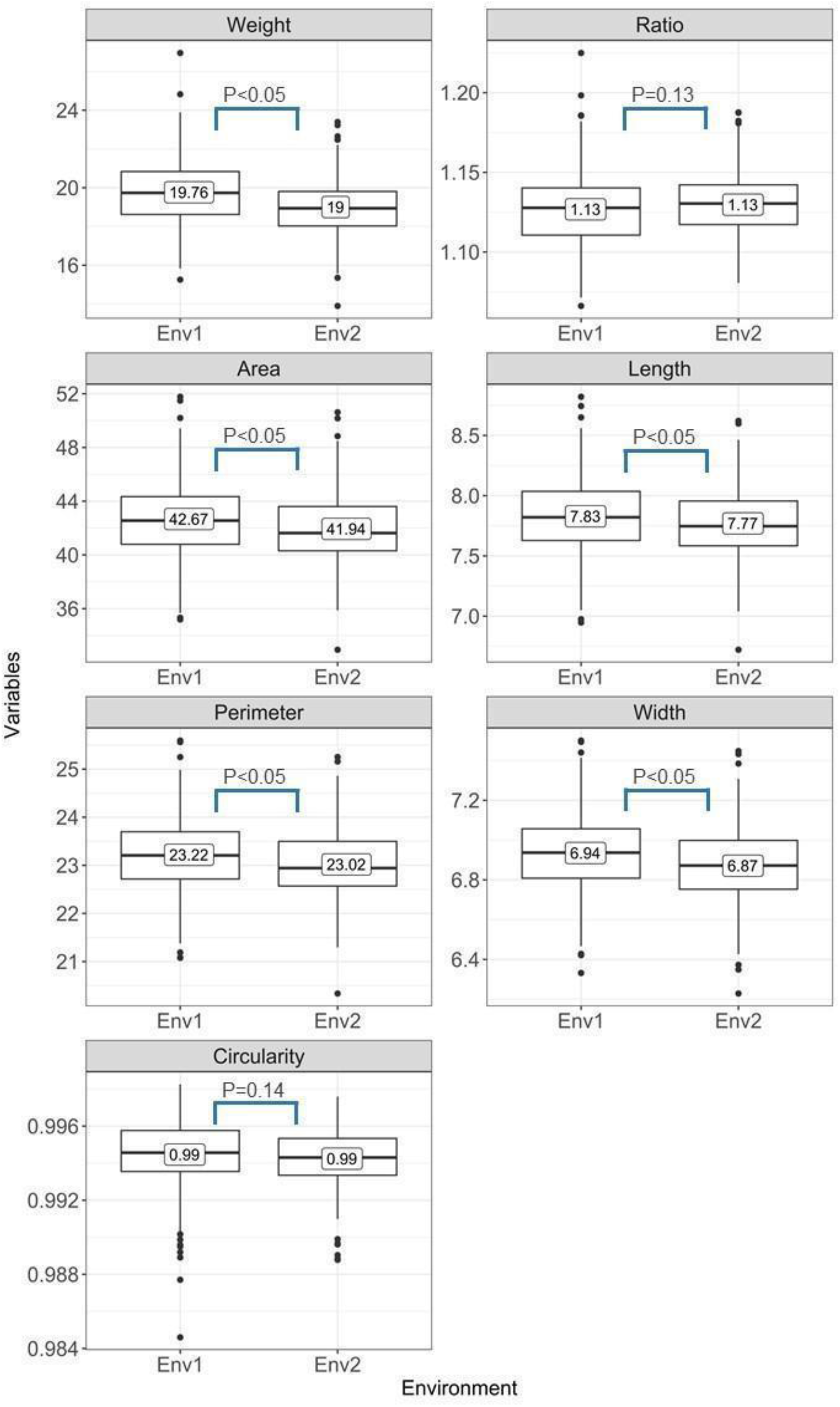
Boxplots of corrected phenotypes for morphological characteristics and hundred-seed weight for environments 1 (Env.1) and 2 (Env.2). P-values less than 0.05 indicate significance at 5% error probability level according to the Student’s t-test. Weight: hundred-seed weight (g); Area: seed area (mm^2^); Perimeter: seed perimeter (mm); Circularity: seed circularity; Ratio: length/width ratio; Length: seed length (mm); Width: seed width (mm); Env.1: with fungicides for Asian soybean rust. Env. 2: without fungicides for Asian soybean rust.

### 2.1. Plant Material

The inbred lines of this study were originated from a partial diallel cross (8×8), conducted by the recurrent selection program 42 of the Sector of Genetics Applied to Autogamous Species (SGAEA) of ESALQ/USP (Rocha et al., 2018; Bornhofen, 2019). The non-transgenic parents were divided into: (i) 8 inbred lines that are tolerant to stem canker (*Diaporthe phaseolorum* f. sp. *meridionalis*), cyst nematode (*Heterodera glycines*), and with a high oil content; and (ii) 8 inbred lines with the same tolerances of (i), and also to Asian rust (*Phakopsora pachyrhizi*). The objective of the crosses was to obtain lines with high agronomic performance, high oil content and tolerance to Asian rust, which is considered the most destructive disease affecting soybean production worldwide (Kashiwa et al., 2020). From each of the 64 crosses, 4 lines were selected in F5:7 with a selection pressure of 33.33%, corresponding to a total of 256 lines for experimentation, 16 parental lines, and three checks (BRS 133, CB 07-958, and IAC 100). In total, we tested 275 soybean genotypes with distinct morphological and agronomic characteristics.

### 2.2 Experimental Design

The experiments were conducted in Piracicaba, São Paulo, Brazil (22°42′15.2″ S and 47°38′24.6″ W, altitude 460m) from November/2020 to March/2021. We employed a Federer’s augmented block experimental design (Federer, 1956), with 259 treatments, composed of 256 experimental inbred lines and 3 checks. A total of 8 blocks with 35 genotypes each (32 inbred lines and 3 checks) was used, considering two replicates, and totalizing 560 experimental units. In addition to the experimental lines, experiments were carried out with the 16 parental lines and the three common checks in a randomized block design with two replicates.

Seed evaluations were performed in two environments: (Env.1) one with total management for controlling rust (spraying of fungicides) and end-of-cycle foliar diseases (fluxapyroxad and pyraclostrobin at a dose of 58.45 + 116.55 g ai.ha^−1^); and (Env.2) another with no management control for Asian soybean rust (control of only end-of-cycle foliar diseases: carbendazim at a dose of 300 g ai. ha^−1^). All management practices adopted in the experiments followed the technical recommendations for soybean cultivation in the region. Harvesting was performed on plants with full maturation using a combined harvester from experimental plots. After drying the seeds at room temperature (~13% moisture), we determined the HSW by measuring the mass of 100 seeds, expressed in grams, randomly sampled from the total produced by each experimental unit.

### 2.3 Image Acquisition

RGB images of 100 seeds from each experimental unit were obtained using an Epson Expression 11000XL Photographic Scanner, with a resolution of 300 dots per inch (dpi) and a size of 257 × 364 mm. The seeds were placed on the top of a 297 × 420 mm transparent acetate sheet using two different configurations (Fig. 1-1d): (Conf.1) the seeds were arranged on the scanner surface in a sparsely distributed manner (10 rows, 10 columns), and (Conf.2) randomly distributed. These different configurations were used in order to evaluate the segmentation process in not only a controlled scenario, but also in a more realistic configuration to provide a new tool for plant breeders. Additionally, a graduation in centimeters was used on the seed sheet together with a scale with RGB colors, for respectively calculating the real dimensions of the seeds and reducing the color detection error depending on the light condition.

### 2.4 Image Segmentation and Characterization

For image processing, we used the Python v.3 programming language (Van Rossum; Drake, 2009) together with the OpenCV library (Pulli et al., 2012). Each image was processed using the same filtering and segmentation process. For segmentation, we employed the following mathematical morphological operations: (i) erosion with a kernel size of 25 pixels (Fig. 1–2a); (ii) closure with a kernel size of 11 pixels (Fig. 1–2b); (iii) Averaging blur filtering with a kernel size of 25 pixels (Fig. 1–2c); (iv) contrast enhancement of 2.5 and brightness of 50 (Fig. 1–2d); and (v) dilation with a kernel size of 25 pixels. To establish a segmentation mask (Fig. .1–2f), we used the Otsu’s binarization (Fig. 1–2e) method, which enabled the definition of connected components using the Grana (Grana et al., 2010) and the Wu (Wu et al., 2009) algorithms.

With a subset of images with sparsely distributed seeds (Conf.1), the components obtained with the segmentation process were used to estimate the distribution of the number of pixels per seed, enabling the establishment of the average number of pixels in a single seed. Based on this *ad-hoc* criterion, we considered the interval of 500 – 8,000 pixels to classify a detected component as a single soybean seed, forming an automatic method to characterize the identified components as seeds or other image elements. Boundary boxes for each seed were calculated using the contour lines of the soybean seeds, enabling a sub-image definition. Each soybean seed was stored as a separate image after its background was removed.

With the image segmented, the best ellipse around the segmented region was estimated (Fig. 1–2i) using the approach of Fitzgibbon and Fisher (1995). We converted the measures of pixels to millimeters by calculating the number of pixels present in 1cm of the graduation image (1 pixel = 0.084 mm). Based on this conversion, we estimated the size and conformation parameters of the seeds: length (L) (major axis of the ellipse); width (W) (minor axis of the ellipse); length/width ratio (LWR) (Equation 1) (Cervantes et al., 2016); area (AS) (Equation 2); perimeter (PS) (Equation 3) (Villarino, 2005); and circularity (CS) (Equation 4) (Rovner and Gyulai, 2007).

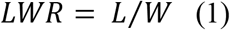

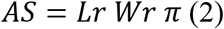

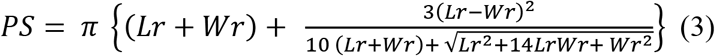

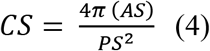

where *Lr* is the radius of the ellipse’s major axis, *Wr* is the radius of the ellipse’s minor axis, and *PS* is an approximation of the perimeter value.

We evaluated the methodology performance considering each environment and seed distribution configuration separately, by calculating: (i) the number of images that showed 100% of the components correctly identified (hits) and without false positives; (ii) the number of images that showed unidentified components (misses); and (iii) the number of images that had components falsely detected compared to the real seed number (false positives). To investigate the causes of incorrect component identification for those images with misses and false positives, we calculated the average of hits, misses, false positives, and the deviation of the real and estimated number of components for each image. We used the following formula for deviation:

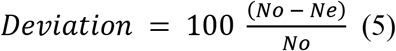

where No and Ne are, respectively, the actual and estimated number of components (Barbedo, 2012).

### 2.5 Segmentation Methodology Validation

To evaluate the performance of the image segmentation methodology in other crops, we also captured seed images of common bean (*Phaseolus vulgaris*), maize (*Zea mays*), and chickpea (*Cicer arietinum*), which are crops with seeds as the main commercial product. RGB images were obtained using the same procedures for capturing images of soybean seeds. Additional adjustments were performed in the segmentation parameters for each crop (Table 1), with the exception of the common bean images, in which the same parameters used in the soybean seeds were used.

**Table 1.** Specifications of segmentation methodologies for each plant species.

| Species | Erosion | Closing | Gaussian blur | Contrast | Brightness | Dilation | Number of pixels per seed* |
| --- | --- | --- | --- | --- | --- | --- | --- |
| *Glycine max* | 25 | 11 | 25 | 2.5 | 50 | 25 | 500 - 8,000 |
| *Phaseolus vulgaris* | 25 | 11 | 25 | 2.5 | 50 | 25 | 500 - 8,000 |
| *Zea mays* | 50 | 31 | 45 | 2.5 | 50 | 50 | 500 - 8,000 |
| *Cicer arietinum* | 25 | 11 | 25 | 2.5 | 20 | 25 | 500 - 9,000 |

\* Minimum and maximum pixel limit per seed.

### 2.6 Statistical Analyses of Phenotypic Data

The analysis of HSW and morphological measure estimates was performed using linear mixed-effects models implemented in the software R (R Core Team, 2021) together with the ASReml-R (Butler et al., 2009) package. We tested the significance of fixed and random effects employing respectively the Wald F test and an analysis of deviance (ANADEV) with likelihood ratio tests (LRTs), considering a Chi-square test. Variance components were estimated via the restricted maximum likelihood method and the best linear unbiased predictor (REML/BLUP) (de Resende, 2000). For the inbred lines’ phenotypic data, we used the following model for each environment:

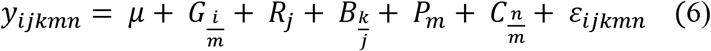

where *y*_*ijkmn*_ is the phenotypic measure of the *i*-th genotype, in the *j*-th repetition, within the *k*-th block and the *m*-th group, and considering the *n*-th check; μ is the overall population mean; *G*_*i*⁄*m*_ is the random effect of the *i*-th genotype within the *m*-th group (Gi ~ N(0, σ^2^_G_), with σ^2^_G_ representing the additive genetic variance); R_j is the effect of the *j*-th repetition; *B*_*k*⁄*j*_ is the random effect of the *k*-th block within the *j*-th repetition; *P*_*m*_ is the fixed effect of groups (inbred lines and checks); *C*_*n*⁄*m*_ is the fixed effect of the *n*-th check within groups; and *ε*_*ijkmn*_ is the random effect of the associated errors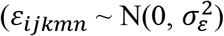, with 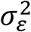 representing the error variance).

We also employed a joint analysis for the two locations, including a fixed effect of the environment (E) and a random effect for the interaction between genotypes and environments (GE), according to the model below:

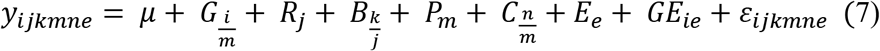

Mixed-effect model equations were also used in the statistical analysis of the parental experiments. The variance components were obtained using the REML/BLUP method, according to the following models for individual and joint analysis of the environments:

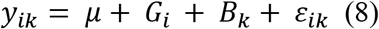

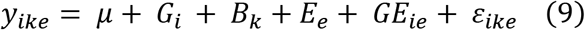

where *y*_*ik*_ is the phenotypic measure of the *i*-th genotype, in the *k*-th block; μ is the overall population mean; *G*_*i*_ is the random effect of the *i*-th genotype (Gi ~ N(0, σ^2^_G_)); *B*_*k*_ is the random effect of the *k*-th block; *E*_*e*_ is the fixed effect of the *e-th* environment; *GE*_*ie*_ is the random effect for the interaction between genotypes and environments; and *ε*_*ike*_ is the random effect of the associated errors 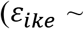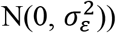

The generalized heritability (broad-sense heritability) for each trait was calculated according to Cullis et al. (2006):

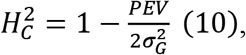

where PEV is the prediction error variance (average variance of individual plant comparisons) and 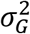 is the genetic variance.

The experimental accuracy was estimated to assess the general quality of the data through the equation:

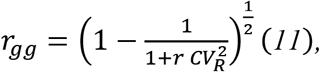

where *CV*_*R*_ = *CV*_*G*_ ⁄*CV*_*ε*_, with 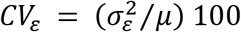,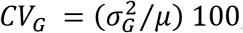, and *μ* is the grand mean (de Resende and Duarte, 2007).

Using the corrected phenotypes of each genotype, Pearson’s correlations were performed between the variables, with significance by means of Student’s t test.

### 2.7 Predictive System Based on Machine Learning: Structured Dataset for Prediction

For predicting the HSW in soybean using the morphological characterization, we employed the ML algorithms: support vector machine for regression (SVR) (Vapnik, 2000), random forest (RF) (Breiman, 2001), multilayer perceptron (MLP) neural network (Pal and Mitra, 1992), and adaptive boosting (AdaBoost) (Freund and Schapire, 1997). For the input data in the prediction system, we employed for each genotype the median value of the 6 morphological characteristics measured for the segmented seeds (L, W, LWR, AS, PS, CS). These medians were first calculated considering the totality of segmentation pictures (100 seeds), and later fractionated at random to use information from only 50, 25, and 10 seeds in the predictions. For evaluation purposes, we employed a k-fold (k=4) cross-validation (CV) technique repeated 50 times using the Python scikit-learn library (Pedregosa et al., 2011). The Pearson’s correlation coefficient (R) and the mean squared error (MSE) were used as evaluation metrics for the regression models, as shown in the equations below:

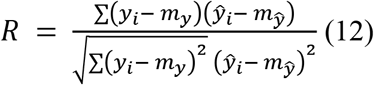

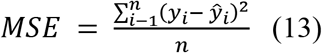

where *y*_*i*_ and *ŷ*_*i*_ are the observed and predicted phenotypes, respectively; *m*_*y*_ is the average of the observed phenotypes; *m*_*ŷ*_ is the average of the predicted phenotypes; and *n* is the total number of samples in the test set.

### 2.8 Predictive System Based on Machine Learning: Image Dataset for Prediction

In addition to the regression models applied to the morphological characteristics, we employed a transfer learning approach, using convolutional layers from CNNs as feature extractors, and the same ML algorithms from the previous section (SVR, RF, MLP, and AdaBoost) for the regression tasks (Fig. 1–3b). To obtain features describing the images, we used the Keras library of Python v. 3 (Chollet et al., 2018) and the following CNN architectures: VGG-16 (Simonyan and Zisserman, 2014), VGG-19 (Simonyan and Zisserman, 2014), ResNet-50 (HE et al., 2016), InceptionV3 (Szegedy et al., 2016), and InceptionResNetV2 (Szegedy et al., 2017). The same evaluation approaches from the previous section were performed for the transfer learning process.

## 3 Results

### 3.1 Segmentation Methodology

The segmentation methodology proposed was mainly based on mathematical morphological operations. Firstly, we used an erosion operation with the aim of disconnecting pixels from different seeds located close to seed boundaries. The inclusion of this operation enabled an efficient distinction of different seeds, even with no significant distancing observed between them. Then, image closing and lighting transformations were applied to prevent an erroneous recognition of the seed hilum as a separate component. An averaging blur was applied as a low-pass filter to reduce the image details, which enabled an easier distinction between the seed and the background of the image. Subsequently, for the definition of a segmentation mask, we employed the Otsu’s thresholding together with an established criterion for seed selection: the exclusion of image components with the quantity of pixels below 500 or above 8,000. Finally, the segmented images were dilated with the same erosion kernel and used to cut the original image for: (i) obtaining the final segmentation; (ii) delimiting the ellipses around the seeds; and (iii) calculating the morphological characteristics. After segmentation, we observed a reduction of ~86.54% in the image size (from 1.76 MB to ~237 KB).

From 596 experimental plots in each environment, a total of 1,150 and 1,190 images were obtained in Env.1 and Env.2, respectively. Out of this total, 1,094 images of Env.1 (547 experimental plots) and 1,186 images of Env.2 (593 experimental plots) were captured for both Conf.1 and Conf.2 seed dispersions. We could verify that from the total of 2,280 images from Env.1 and Env.2, 1,999 images (~87.67%) showed the 100 components correctly identified and without false positives (Table 2). There were only 204 images (~8.95%) with unidentified components and 77 (~3.38%) with falsely detected components. Although the exclusion of components with the quantity of pixels outside the interval of 500 – 8,000 was carried out, the main reason for the misidentification of image components as seeds was the presence of the graduation in centimeters and scale with RGB colors next to the seeds, which in the minority of cases was considered a component.

**Table 2.** Segmentation evaluation of the proposed approach.

|  | Environment | Conf.1* | Conf.2* | Total |
| --- | --- | --- | --- | --- |
| Images with 100% hits and no false positives | 1 | 507 (~92.69%) | 437 (~79.89%) | 944 (~86.29%) |
|  | 2 | 583 (~98.31%) | 472 (~79.60%) | 1055 (~88.95%) |
| Images with misses | 1 | 6 (~1.10%) | 75 (~13.71%) | 81 (~7.40%) |
|  | 2 | 7 (~1.18%) | 116 (~19.56%) | 123 (~10.37%) |
| Images with false positives | 1 | 34 (~6.22%) | 35 (~6.40%) | 69 (~6.31%) |
|  | 2 | 3 (~1%) | 5 (~1%) | 8 (~1%) |

\*Conf.1: 100 seeds sparsely distributed on the scanner; Conf.2: 100 seeds densely distributed on the scanner. Environment (Env.) 1: with fungicides for Asian soybean rust. Env. 2: without fungicides for Asian soybean rust.

The quantity of images with incorrect component identifications (misses or false positive) was 150 in Env.1 and 131 in Env.2, with a high average of hits (~99.18%) together with a low average of false positives (~0.41%), culminating in a total average deviation from the real and the estimated number of components of only ~0.81% (Table 3).

**Table 3.** Segmentation evaluation of the proposed approach considering the images with any incorrect component identification. The hits, misses and false positives columns indicate the average number of correctly identified, unidentified and falsely identified components present in an image, respectively. The deviation column reveals the average percent error in component identification.

| Environment | Configuration* | Hits<br>(Mean) | Misses<br>(Mean) | False<br>Positives<br>(Mean) | Deviation (%)<br>(Mean) |
| --- | --- | --- | --- | --- | --- |
| 1 | 1 | 99.85 | 0.15 | 0.93 | 0.15 |
|  | 2 | 99.05 | 0.96 | 0.38 | 0.95 |
| 2 | 1 | 99.20 | 0.80 | 0.30 | 0.80 |
|  | 2 | 98.64 | 1.36 | 0.04 | 1.36 |

\*Configuration 1: 100 seeds sparsely distributed on the scanner; Configuration 2: 100 seeds densely distributed on the scanner. Env.1: with fungicides for Asian soybean rust. Env. 2: without fungicides for Asian soybean rust.

We could observe that ~79.73% of the images obtained in Conf.2 showed all components correctly identified, as opposed to ~95.61% of images in Conf.1 (Table 2). Even with such a difference, when we analyzed the images with identification errors, we observed that the average of hits in Conf.1 was ~99.53% and in Conf.2 was ~98.85%, representing small differences with an error rate of ~0.68% (Table 3). Even with eventual errors, all images had more than 98% of the seeds correctly identified, showing that even in non-ideal seed dispersions, the methodology is capable of isolating seeds.

### 3.2 Methodology Evaluation with Images of Other Plant Species

Through the evaluation of the image segmentation methodology in seed images of other crops, we observed that the developed workflow also resulted in an efficient seed segmentation, regardless of the size, shape, texture, or color of the seed (Fig. 2). It is worth mentioning that for *Cicer arietinum* and *Zea mays* it was necessary to perform small adjustments in the image segmentation parameters (Table 1), due to the discrepant seed morphologies in comparison to soybean.

For *Phaseolus vulgaris* and *Cicer arietinum* seeds, the methodology developed correctly detected the 100 components (Table 4). For *Zea mays*, there was only one unidentified component in both images with sparsely (Conf.1) and densely (Conf.2) distributed seeds. In the images of the three species, there were no falsely detected components compared to the real number (false positives). These results indicate the potential generalization capacity of the proposed segmentation methodology.

**Table 4.** Evaluation of the segmentation methodology for other plant species. The hits, misses and false positives columns indicate the number of correctly identified, unidentified and falsely identified components present in an image, respectively. The deviation column reveals the percentage of error in component identification.

| Species | Configuration* | Hits | Misses | False Pos. | Deviation (%) |
| --- | --- | --- | --- | --- | --- |
| *Phaseolus vulgaris* | 1 | 100 | 0 | 0 | 0 |
|  | 2 | 100 | 0 | 0 | 0 |
| *Zea mays* | 1 | 99 | 1 | 0 | 1 |
|  | 2 | 99 | 1 | 0 | 1 |
| *Cicer arietinum* | 1 | 100 | 0 | 0 | 0 |
|  | 2 | 100 | 0 | 0 | 0 |

\* Configuration 1: 100 seeds sparsely placed; configuration 2: 100 seeds densely placed.

### 3.3 Phenotypic Data Analysis

Six morphological characteristics of 234,000 seeds were measured for individual seeds of 2,340 images from 275 soybean genotypes. We observed that the distribution of these phenotypes from image analysis followed a normal distribution with continuous variation (Fig. 5). From the estimated mixed-effects models for such characteristics (Conf.1 and Conf.2), we observed significant genotype effects (P<0.0001) in the models created for both the inbred and parental lines (Supplementary Tables 2-3), which indicates that the methodology of segmentation and morphological characterization of the images was able to capture the genotypic variability in the lines.

**Figure 4.**
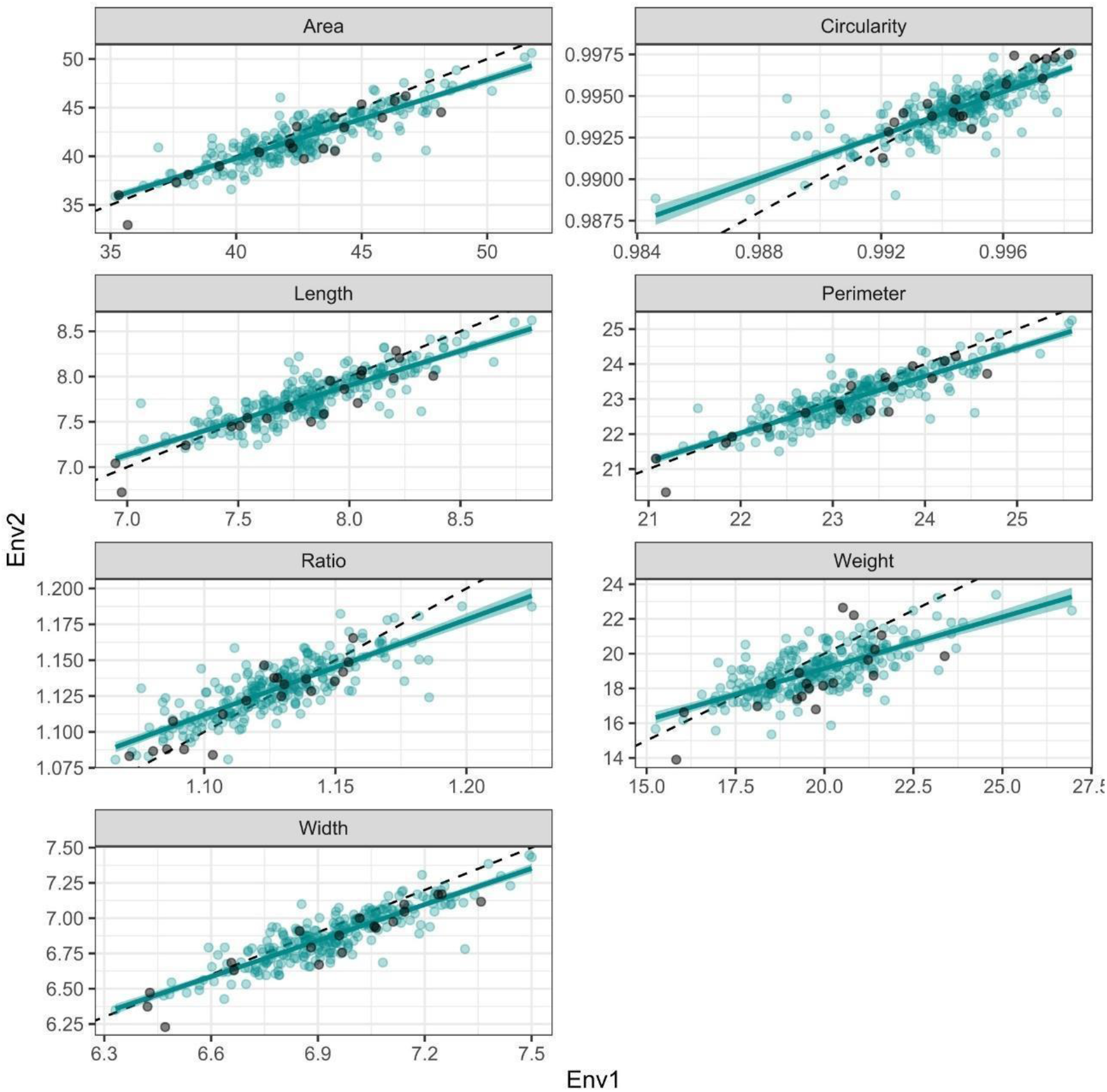
Scatter plots comparing the performances of the genotypes evaluated under two stress conditions for hundred-seed weight and six seed morphological traits. Blue dots: Inbred lines. Gray dots: Parental lines. The dots in the lower right triangle (proportion of the point cloud located below the 45° diagonal) indicate individuals with higher values of variables in the environment with rust control. Weight: hundred-seed weight (g); Area: seed area (mm^2^); Perimeter: seed perimeter (mm); Circularity: seed circularity; Ratio: length/width ratio; Length: seed length (mm); Width: seed width (mm). Env.1: with fungicides for Asian soybean rust. Env. 2: without fungicides for Asian soybean rust.

**Figure 5.**
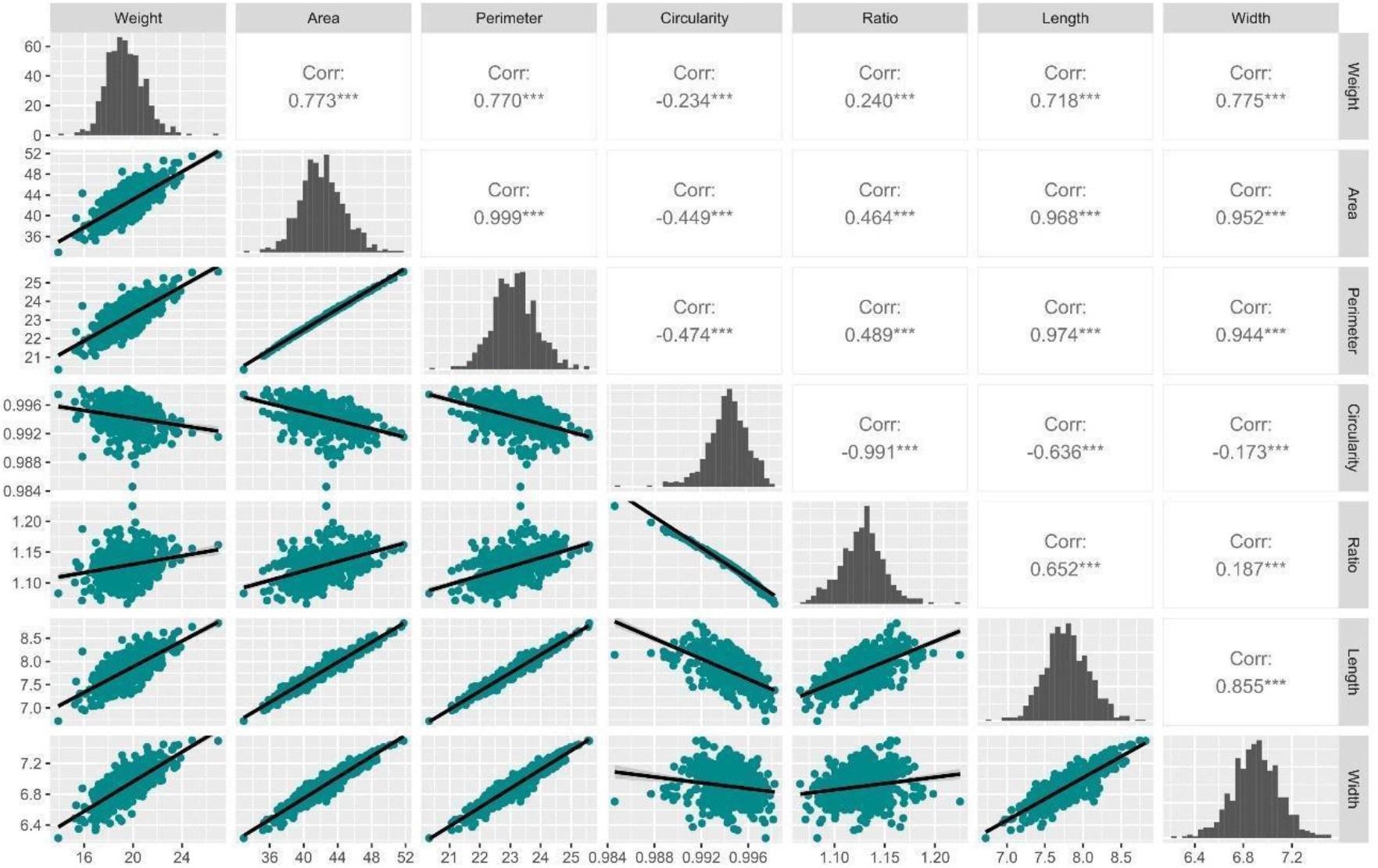
Scatter plots of the correlation variables for the experiments. The distribution of each variable is shown on the diagonal. On the bottom of the diagonal: the bivariate scatter plots with a fitted line are displayed. On the top of the diagonal: the value of the correlation plus the significance level as stars. ‘***’, ‘**’, ‘*’, ‘.’ Significant at the 0.0001, 0.001, 0.01, 0.05 probability levels, respectively. Weight: hundred-seed weight (g); Area: seed area (mm^2^); Perimeter: seed perimeter (mm); Circularity: seed circularity; Ratio: length/width ratio; Length: seed length (mm); Width: seed width (mm).

In general, estimates of heritability in the broad sense were high, ranging from 74% to 95%, for HSW and length/width ratio, respectively (Supplementary Tables 2-3). Interestingly, the HSW ranged from 13.90 to 26.95 g, the seed area from 32.94 to 51.77 mm^2^, the perimeter from 20.34 to 25.59 mm, the length from 6.72 to 8.82 mm and width from 6.23 to 7.50 mm, respectively for the same genotypes: IAC 100 (control susceptible to *P. pachyrhizi*) and the USP 17-42.244 line. Such results together with the high correlations observed between the characteristics evaluated (morphological characteristics and hundred-seed weight) (Fig. 5) demonstrate the reliability of the segmentation methodology, mainly because of the real strong relatedness observed between these parameters in a soybean breeding evaluation (Niu et al., 2013).

We observed strong negative correlations between the characteristics of circularity and length/width ratio (−0.99), suggesting that these parameters are inversely proportional. Seed circularity ranged from 0.98 to 1.00 for the inbred lines USP 17-42.361 and USP 17-42.022, respectively, and the length/width ratio from 1.07 to 1.22 for the same lines, but reversed (USP 17-42.022 and USP 17-42.361, respectively).

Seed circularity is a measure of how closely the seed resembles a circle (Cervantes et al., 2016). The values can vary from zero to one, with one corresponding to a seed with the appearance of a perfect circle (Cervantes et al., 2016). In general, the seeds showed a high circularity (~1), which is confirmed by the values of the length/width ratio close to one and below 1.50, which refers to the maximum limit defined by Balkaya and Odabas (2002) for the seed to be classified as round. Additionally, it was observed that the circularity had a negative correlation with all other characteristics. It is noteworthy that seed length is highly correlated negatively with circularity (−0.64) and positively with the length/width ratio (0.65), suggesting that seed shape may depend on seed length.

Considering the checks IAC 100 and BRS 133 as susceptibility controls and CB 07-958 as a tolerance control for *P. pachyrhizi*, we observed the smallest means of area, perimeter, length, and width of the seeds for the checks IAC 100 and BRS 133, in comparison with the check CB 07-958 and the other genotypes, demonstrating the highest values of seed shape variables for the genotypes that underwent selections for disease tolerance. Regarding the HSW, only the IAC 100 had lower values (14.87 g), compared to the BRS 133 (18.36 g), CB 07-958 (18.31 g), and other genotypes (19.41 g). For circularity and length/width ratio, the controls showed very similar values compared to the overall mean of the inbred lines and parental lines.

We observed that the differences between the corrected phenotypes from the models created for Env.1 and Env.2 for HSW, area, perimeter, length, and width were significant by the Student’s t-test, with a slight reduction in the values of these characteristics in Env.2 (Fig. 3). Considering the joint model for Env.1 and Env.2, we also observed significant effects of the interaction between genotypes and environments only for the length, circularity, and length/width ratio of the seed for the experiments with parental lines (Supplementary Table 2), indicating that there was a differential performance in the parents across environments.

In addition to the statistical tests, the same findings were observed in the scatter plots created for contrasting Env.1 and Env.2 (Fig. 4). In descending order, the proportion of point clouds located below the 45° diagonal line (perfect correlation) was 76.28% for width, 75.91% for HSW, 72.99% for seed area, 72.99% for perimeter, 67.52% for length, and 63.5% for circularity (Table 5), indicating a majority of genotypes with superior performance in the environment with Asian soybean rust control (Env.1). The length/width ratio was the only variable with superior performance in the presence of rust (Env.2), with 62.04% of the points located above the 45° line, because it is inversely proportional to the circularity.

**Table 5.** Proportion of the point cloud located above the 45° diagonal (variable in environment 2 was higher than in environment 1: Env 2>Env 1). Proportion of the point cloud located below the 45° diagonal (variable in environment 1 was higher than in environment 2, Env 1>Env 2). Correlation between the variable in environment 1 and environment 2.

| Variable | Env 2>Env 1(%) | Env 1>Env 2 (%) | Correlation |
| --- | --- | --- | --- |
| HSW | 24.09 | 75.91 | 0.70 *** |
| AS | 27.01 | 72.99 | 0.87 *** |
| PS | 27.01 | 72.99 | 0.87 *** |
| L | 32.48 | 67.52 | 0.86 * |
| W | 23.72 | 76.28 | 0.88 *** |
| CS | 36.50 | 63.50 | 0.80 ns |
| LWR | 62.04 | 37.96 | 0.82 ns |

HSW: hundred-seed weight; AS: seed area; PS: seed perimeter; L: seed length; W: seed width; CS: seed circularity; LWR: length/width ratio. Env.1: with fungicides for Asian soybean rust. Env. 2: without fungicides for Asian soybean rust. '\*\*\*', '\*\*', '\*' Significant at the 0.001, 0.01, 0.05 probability levels, respectively.

### 3.4 Performance Metrics of Machine Learning Models

In this study, we tested the performance of ML models to predict the HSW from the image-based morphological measurements of the seeds. We observed that using the size and conformation parameters calculated based on the images containing 100 seeds, model performances achieved a mean predictive ability of 0.71 and a mean squared error of 3.15 (Fig. 6).

**Figure 6.**
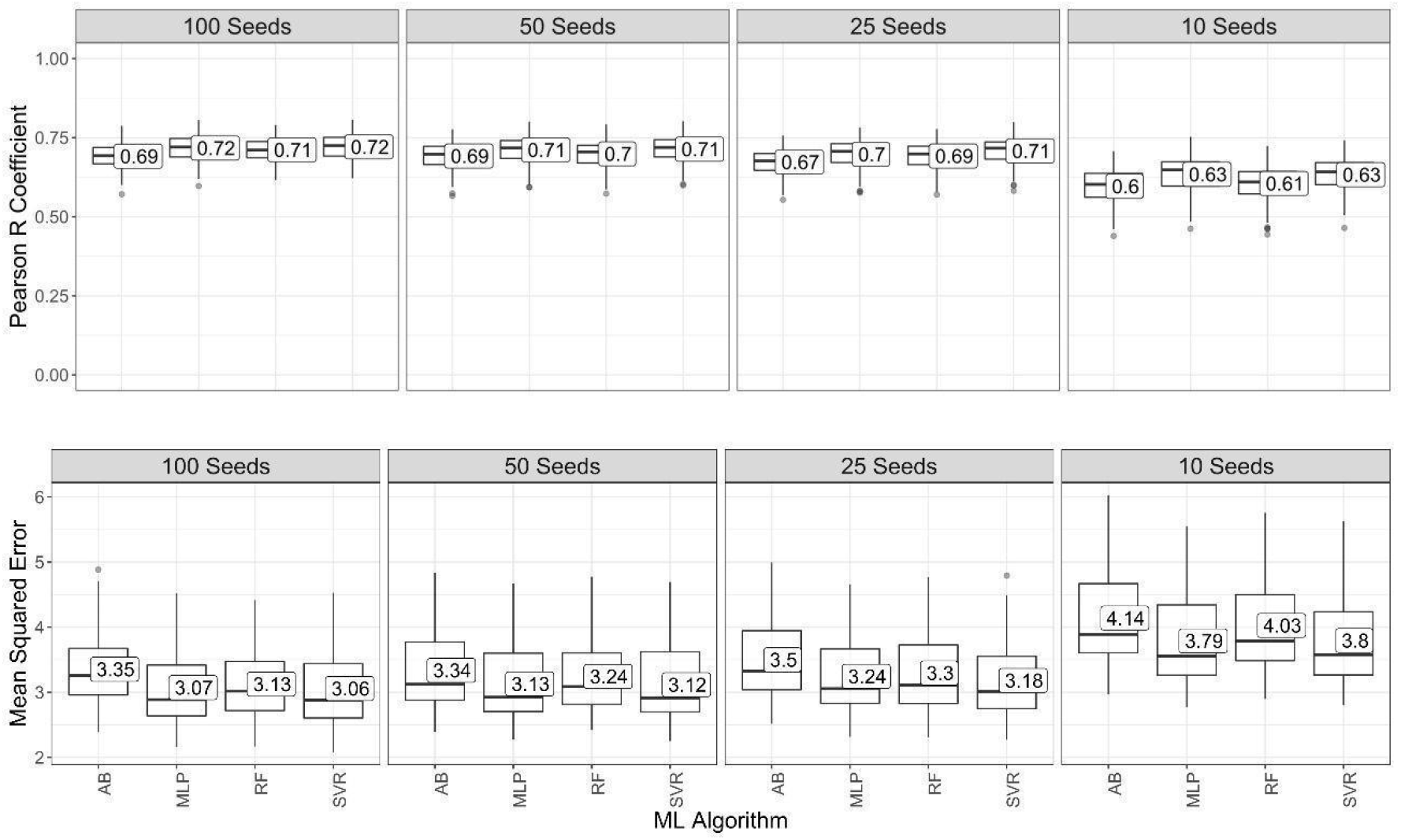
Predictive performance evaluation of machine learning (ML) algorithms (*AdaBoost, multilayer perceptron* (*MLP*), *random forest* (*RF*), and *support vector regression* (*SVR*)) for prediction of the hundred-seed weights using the images of the two environments considering the Pearson R coefficient and the mean squared error as evaluation metrics.

Furthermore, we evaluated the model performances using measurements from images with smaller numbers of seeds, aiming to check the feasibility of reduced datasets for this prediction task. When using the morphological characteristics of 50 and 25 seeds for the prediction, we observed a small reduction in the mean predictive ability of the models to 0.70 and 0.69, respectively. With the use of 10 seeds, a larger reduction in the predictive ability was observed, together with a significant increase in the predictive errors (~25%).

The AdaBoost, MLP, RF, and SVR models showed similar results in the predictions. It was possible to verify that the predictions made with seed data from the environment with total disease control (Supplementary Fig.1) showed higher accuracies and lower mean squared errors compared to those from the environment without control of Asian soybean rust (Supplementary Fig. 2), possibly due to the lower influence of the environment on the variability of seed morphology.

We performed a comparative analysis among the combination of CNN architectures for feature extraction and ML algorithms. In a comprehensive comparison among all architectures for feature extraction, ResNet-50 was the most effective with a mean accuracy of 0.60 (Figure 7). Among the transfer learning models, the algorithm based on ResNet-50 features with MLP (ResNet-50 + MLP) obtained the best performance (Table 6).

**Table 6.** Tukey’s HSD test based on predictive performance evaluation of the combination of convolutional neural network (CNN) architectures for feature extraction and machine learning (ML) algorithms (*AdaBoost*, *multilayer perceptron* (*MLP*), *random forest* (*RF*), and *support vector regression* (*SVR*)) for prediction of the hundred-seed weights using the images of the two environments considering the Pearson R coefficient and the mean squared error as evaluation metrics.

| Algorithms | Accuracy |  | MSE |  |
| --- | --- | --- | --- | --- |
| ResNet-50/MLP | 0.64 | a | 4.00 | f |
| ResNet-50/SVR | 0.61 | ab | 4.10 | ef |
| VGG-16/SVR | 0.61 | ab | 4.20 | def |
| ResNet-50/RF | 0.59 | ab | 4.12 | def |
| VGG-19/SVR | 0.57 | abc | 4.38 | cdef |
| VGG-19/RF | 0.57 | abc | 4.32 | def |
| VGG-16/RF | 0.57 | abcd | 4.41 | cdef |
| ResNet-50/AdaBoost | 0.56 | abcde | 4.40 | cdef |
| InceptionV3/MLP | 0.55 | abcde | 4.84 | cdef |
| VGG-16/MLP | 0.52 | bcdef | 5.44 | bcde |
| InceptionV3/RF | 0.51 | bcdef | 4.65 | cdef |
| VGG-19/AdaBoost | 0.49 | cdef | 4.82 | cdef |
| VGG-16/AdaBoost | 0.49 | cdef | 4.81 | cdef |
| InceptionV3/SVR | 0.47 | def | 4.99 | cdef |
| InceptionV3/AdaBoost | 0.46 | efg | 5.15 | cdef |
| VGG-19/MLP | 0.42 | fgh | 7.01 | a |
| InceptionResNetV2/MLP | 0.37 | ghi | 6.64 | ab |
| InceptionResNetV2/RF | 0.35 | hi | 5.52 | bcd |
| InceptionResNetV2/SVR | 0.33 | hi | 5.73 | abc |
| InceptionResNetV2/AdaBoost | 0.32 | i | 5.79 | abc |

Means followed by different letters are statistically different at 0.05 probability level.

**Figure 7.**
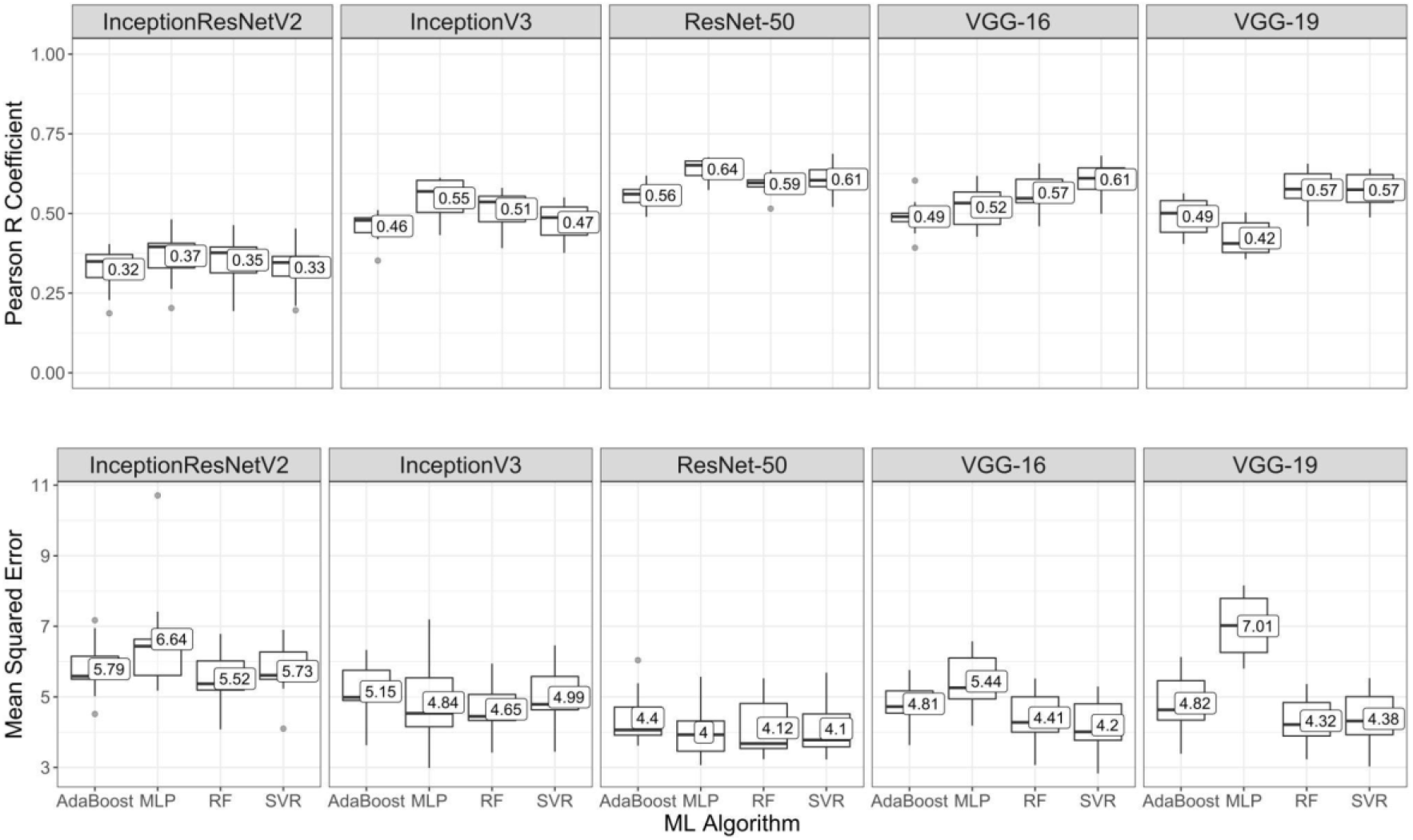
Predictive performance evaluation of machine learning (ML) algorithms (*AdaBoost, multilayer perceptron* (*MLP*), *random forest* (*RF*), and *support vector regression* (*SVR*)) for prediction of the hundred-seed weights using the images of the two environments considering the Pearson R coefficient and the mean squared error as evaluation metrics.

The difference in the mean accuracy of prediction of the HSW in soybean between models using quantitative data from morphological characterization and transfer learning approaches was only 0.18. We observed that predictions based on deep learning features performed similarly to those ML models using morphological seed data, which corroborates the efficiency of morphological measures as feature extractors.

## 4 Discussion

### 4.1 Seed segmentation pipeline

In the last few decades, image analysis software programs for seed evaluation have emerged with the use of well-established image processing methodologies, allowing the evaluation of several morphological characteristics, which have been little explored due to the difficulty of manual measurements, such as perimeter and circularity (Lamprecht et al., 2007; Igathinathane et al., 2008; Tanabata et al., 2012; Gehan et al., 2017). However, such approaches have shown diverse limitations, especially for segmenting seeds under inconsistent lighting conditions or that are densely arranged and have physical contact with each other (Yang et al., 2021). When seeds are closely placed, they are often detected as a unified image component, implying the misrecognition of isolated seeds (Toda et al., 2020). Therefore, there is a need for new image segmentation approaches, in which the laborious process of sparsely disposing the seeds without overlaps is not necessary. Bypassing such limitations, we developed an analysis framework with a high potential to be expanded into an automated breeding practice, optimizing seed phenotyping and allowing the investigation of many samples in a faster, more accurate, and less expensive way.

In our study, we were able to measure and analyze ~234,000 soybean seeds of 275 genotypes, correctly identifying more than 98% of the seeds present in the images in all tested cases (Conf.1 and Conf.2) (Tables 2-3). Taking into account the reduced time for image acquisition in Conf.2 and the small differences in error rate (0.68%) between both configurations, we consider that the use of the proposed methodology together with the acquisition of images with densely distributed seeds can be advantageous for seed morphology evaluation in breeding programs, assisting breeder’s decisions.

Even with such importance, little efforts to solve this problem have been reported in the literature. Although Yang et al. (2021) proposed a deep learning-based method for image segmentation in seeds arranged with physical contact, the authors report a high computational cost and the need of a large amount of data to train the model, hindering the application in smaller datasets. Our image segmentation framework proposed here, on the other hand, is entirely based on mathematical morphological operations, which resulted in an easy implementation pipeline with low computational costs.

Additionally, the application of our pipeline is not restricted to soybean, as demonstrated with seed images from common bean, chickpea and maize. High-throughput seed phenotyping methodologies, such as the framework proposed in our study, represent a promising tool in the development of breeding approaches to ensure food and nutrition security globally, especially for crops whose grain is the main commercial product and for orphan crops, such as chickpea, whose yield increment is still not enough to meet the food demand of a growing world population (Dawson et al., 2019).

### 4.2 Seed morphology evaluation

HTP of seeds can provide useful information for breeding by allowing genotypes to be evaluated in a reduced time with high repeatability. In our study, even employing full-sibs and half-sibs derived from a partial diallel scheme, we observed that small differences in morphological characteristics could be detected. This fact corroborates the importance of these techniques, which can accurately measure a large number of genotypes, unlike the classical measurement techniques (Baek et al., 2021). In addition, morphological characterization of seeds can be very useful to describe soybean cultivars and germplasm accessions (Nelson and Wang, 1989; Baek et al., 2021). HTP directly contributes to more accurate heritability calculations, helping to efficiently estimate the genetic variance and making decision support systems more robust (Araus et al., 2018). Thus, it was observed that the heritability of the HSW was lower than those observed for the morphological characteristics (Supplementary Table 2 and 3). In agreement with these results, the experimental accuracy of the variables was classified as very high, except for the HSW classified as high according to the scale proposed by de Resende and Duarte (2007) (Supplementary Tables 2 and 3).

The contribution of GxE interactions in the variation of traits is crucial in defining breeding goals and strategies for the development of cultivars with high yield potential, as well as stability under different environmental conditions (Marrano and Moyers, 2022). Tolerance to different stresses can be assessed not only through yield, but also via contrasts between yield components. In this scenario, the HSW is an interesting characteristic, due to the high heritability of the trait (Pathan et al., 2013; Kato et al., 2014), as also indicated by our study. However, we observed that the biotic stress caused by the rust severity was not enough to cause major changes in the HSW and morphological characteristics of the seeds. This fact can be explained by the selection process for tolerance to the *P. pachyrhizi* fungus performed in the germplasm used.

Even though we restricted our study to the calculation of 6 morphological characteristics, many other seed measurements can be estimated with our method, due to the flexibility of the pipeline proposed. All the scripts developed are available for use and can be adapted for specific research workflows. Regarding the morphological characteristics evaluated, we confirmed their quantitative nature through Normal distribution fits and continuous variations, which is in agreement with the identification of many quantitative trait locus (QTLs) involved in their genetic control (Hu et al., 2013; Niu et al., 2013; Salas et al., 2016; Baek et al. 2021). Seed conformation characteristics (circularity and length/width ratio) showed different distribution patterns when compared to seed size characteristics (area, perimeter, width and length), as observed in the low correlations between these two classes of parameters (Fig. 5). This result is in agreement with Hu et al. (2013), who point to the genetic independence between seed size and conformation.

High and significant correlation values of the seed size parameters (area, perimeter, width and length) with HSW (Fig. 5) can be observed, which motivated us to use them together with those of seed conformation (circularity and length/width ratio) as predictors of HSW in machine learning models. Considering the high correlations and greater heritabilities of the traits obtained by image processing, we suggest that it is possible to achieve indirect gains in HSW by selecting the seed morphology, taking advantage of a less laborious measurement coupled with an automated image capture system.

### 4.3 Image-based prediction

Our study reinforces the advantages involved in the integration of seed images with proper analytical tools in a plant breeding process. We present a proposal for the HSW prediction based on the morphological data of the seeds obtained from the images, also evaluating its performance with the reduction of the number of seeds per image. We verified that it is possible to use images containing less than one hundred seeds, since the prediction based on the morphological characteristics of 50 and 25 seeds reduced the accuracy by only 0.75% and 1.75%, respectively (Fig. 6). On the other hand, it is important to point out that if there was a gradual decrease in the accuracy of predictions with fewer seeds in the images, presumably with a greater number of seeds, improvements in predictive accuracies may be observed.

Among the approaches for extracting image attributes, transfer learning is an effective method for knowledge adaptation, in which convolutional layers are used as extractors of fixed features (Altuntas et al., 2019; Salaken et al., 2019). In these cases, the last layers of the pre-trained CNN can be coupled with a different machine learning approach (Sravan et al., 2021). These combinations of CNN architectures and conventional learning algorithms make it possible to improve the performance of the resulting model for a specific application (Sravan et al., 2021), as well as creating a predictive framework with greater potential for use.

In this way, we also evaluated the use of CNNs as feature extractors instead of the morphological characterization performed. CNNs are known to provide cutting-edge performance compared to traditional approaches for almost all computer vision tasks, demonstrating a great potential for improving image analysis performance in plant phenotyping (Jiang and Li, 2020). However, when coupling such CNN features with the same ML algorithms, we obtained similar accuracies in the seed morphology-based models, demonstrating the accuracy of such measures. Since CNNs have a deep architecture, and this architecture leads to an elevated complexity with high computational costs, we suggest that the morphological measures can be an efficient alternative for soybean phenotyping.

### 4.4 Application of the study in plant breeding

In this research, we demonstrate the possibility of not only obtaining morphological characteristics of the seeds in a feasible way, but also of predicting an important component of production for the soybean crop, through a workflow for obtaining seed images faster, with less chance of errors and low associated costs. This is a valuable tool for genotype screening, which is a very important process in the early stages of breeding. Generally, in the first generations of plant breeding, selection is based on a very small amount of phenotypic information from individuals, due to the high cost or the impossibility of performing an accurate phenotypic assessment (Rincent et al., 2018). This “phenotyping bottleneck” limits the understanding of how expressed phenotypes correlate with genetic factors and adjacent environmental conditions, slowing the progress on important breeding challenges (Großkinsky et al., 2015). HTP techniques, such as the one proposed in this study, have the potential to accelerate the breeding process, especially in the first generations, when breeders need to select from thousands to millions of individuals those that have the desirable traits (Rincent et al. al., 2018; Watt et al., 2020).

Additionally, despite significant advances in high-throughput genotyping, it can still be difficult and expensive to genotype all individuals for the application of genomic selection, especially for breeding programs with few financial resources (Rincent et al., 2018). HTP, even with a lower predictive capacity, can be attractive to plant breeding, as it requires lower infrastructure investments and is flexible, allowing the workflow to be configured in different locations where breeding programs are performed (Zhu et al., 2021). Thus, by adopting these measures, it is still possible to concentrate financial investments for genotyping in final and more refined stages of improvement.

Despite recent advances in genomics, the lack of adequate phenotyping data has led to poor results in gene/QTL discovery, limiting progress even in genomics-assisted breeding programs (Mir et al., 2019). It is worth mentioning that studies such as this one on seed phenotyping are still very useful not only for identifying QTLs associated with morphological characteristics of seeds, but also for genomic selection approaches that could complement phenomic selection (Zhu et al., 2021).

Given the above, our work has the potential to help future research and the industry to develop automated phenotyping tools, incorporating the proposed analytical workflows. However, the limitation for the use of the pipeline proposed in this study is when the time for weighing the seeds and obtaining the morphological measurements is shorter than the time of image capture and subsequent data processing. The solution to this challenge comes from increasing the accuracy and yield of operations concomitantly with the reduction of costs resulting from the decrease in manpower, which is called economic phenotyping (Mir et al., 2019). Automation and data integration can be considered the main factors responsible for reducing manpower (Mir et al., 2019). The entire process can be automated, including the generation and compilation of phenotypic data (Li et al., 2021). As suggested by Zhu et al. (2021), if there is a combination of real-time image analysis aided by cloud-based approaches, the prediction and subsequent selection of superior genotypes can occur soon after the images are captured.

In the literature, studies on automated seed phenotyping are already found, such as those by Jahnke et al. (2016), who proposed a robot system for automated handling and phenotyping of individual seeds, but their methodology requires a more sophisticated knowledge of robotics, which would limit its wide use. On the other hand, Komyshev et al. (2017) present a method for the automated evaluation of phenotypic parameters of seeds through images captured by mobile devices, but there are important challenges in image processing arising from the interference of irregular lighting captured in the optical sensors of smartphones. Thus, aiming at optimizing the acquisition of images, we suggest for future works the aid of robotics for the development of simple automated instruments designed to increase the speed of obtaining standardized images of the seeds, technologies that are already consolidated in the seed processing sector, but still incipient in improvement pipelines. Thus, the advancement of techniques such as these has the potential to ensure the continuous and efficient development of new soybean cultivars in breeding programs that aim at long-term genetic gain.

## 5 Conclusion

This study presents methodologies for obtaining and studying the morphological characteristics of soybean seeds through a faster workflow for obtaining seed images, with fewer errors and low cost. We also propose HSW predictions based on ML never before reported in the literature. Through the comparative performance between models using quantitative morphological characterization data and transfer learning approaches that use images as input, we show the efficiency of morphological measures as extractors of image features. Our results indicate that ML techniques can be easily adapted to precision tasks in plant phenology, helping to improve current and future breeding programs. The implementation of practical solutions such as this one is valuable to enable the integration of contemporary phenotyping techniques in the workflow for soybean plant breeding.

## Supporting information

Supplementary Results

## Author Contributions

MM and AA performed all analyses, wrote the manuscript, and conceptualized the idea. MM and JB were responsible for the field experiments and soybean phenotyping. All authors contributed to the article and approved the submitted version.

## Acknowledgments

We would like to acknowledge the Fundação de Amparo à Pesquisa do Estado de São Paulo (FAPESP), the Conselho Nacional de Desenvolvimento Científico e Tecnológico (CNPq), and the Coordenação de Aperfeiçoamento de Pessoal de Nível Superior (CAPES).

## SUPPLEMENTARY FILES

Supplementary File 1. Python codes to exemplify the segmentation steps.

Supplementary File 2. Python codes for the segmentation workflow established.

## SUPPLEMENTARY TABLES

Supplementary Table 1. Analysis of deviance (ANADEV) for the hundred-seed weight and morphological characteristics of seeds, in each environment and in the joint analysis of the two environments. Samples of 100 seeds were obtained in experiments containing 256 inbred lines and 3 checks designed in Federer’s augmented blocks.

Supplementary Table 2. Analysis of deviance (ANADEV) for the hundred-seed weight and morphological characteristics of seeds, in each environment and in the joint analysis of the two environments. Samples of 100 seeds were obtained in experiments containing 16 parental lines and 3 checks designed in randomized blocks.

## SUPPLEMENTARY FIGURES

Supplementary Fig. 1. Predictive performance evaluation of machine learning (ML) algorithms (*AdaBoost, multilayer perceptron* (*MLP*), *random forest* (*RF*), and *support vector regression* (*SVR*)) for prediction of the hundred-seed weights using the images of the environment 1 considering the Pearson R coefficient and the mean squared error as evaluation metrics.

Supplementary Fig. 2. Predictive performance evaluation of machine learning (ML) algorithms (AdaBoost, multilayer perceptron (MLP), random forest (RF), and support vector regression (SVR)) for prediction of the hundred-seed weights using the images of the environment 2 considering the Pearson R coefficient and the mean squared error as evaluation metrics.

