## Supplementary Results for "A novel image-based approach for soybean seed phenotyping using machine learning techniques"

**SUPPLEMENTARY TABLES**

Supplementary Table 1. Analysis of deviance (ANADEV) for the hundred-seed weight and morphological characteristics of seeds, in each environment and in the joint analysis of the two environments. Samples of 100 seeds were obtained in experiments containing 256 inbred lines and 3 checks designed in Federer's augmented blocks.

| Env. | Effects Random | | HSW | | L | | W | | AS | | PS | | CS | | LWR | |
|---|---|---|---|---|---|---|---|---|---|---|---|---|---|---|---|---|
| E1 | Inbred lines/G | Variance Component | 3.56 | | 0.10 | | 0.04 | | 8.33 | | 0.63 | | 4.33E-06 | | 6.67E-04 | |
| | | LRT | 124.35 | *** | 287.79 | *** | 242.79 | *** | 284.51 | *** | 2.99 | * | 6641.7 | *** | 229.04 | *** |
| | Block/Rep | Variance Component | 0.17 | | 7.33E-04 | | 2.26E-03 | | 0.29 | | 0.02 | | 2.53E-08 | | 2.49E-07 | |
| | | LRT | 3.66 | * | -0.28 | | 5.80 | ** | 3.72 | * | 278.77 | *** | 6365.9 | *** | 0.00 | |
| | Residual | Variance Component | 2.21 | | 0.02 | | 0.01 | | 1.81 | | 0.14 | | 1.16E-06 | | 1.70E-04 | |
| | | h$^2$ | 0.79 | | 0.91 | | 0.89 | | 0.90 | | 0.90 | | 0.88 | | 0.89 | |
| | Mean | | 19.74 | | 7.83 | | 6.93 | | 42.66 | | 23.21 | | 0.99 | | 1.13 | |
| | Accuracy | | 0.87 | | 0.95 | | 0.94 | | 0.95 | | 0.95 | | 0.94 | | 0.94 | |
| E2 | Inbred lines/G | Variance Component | 2.53 | | 0.09 | | 0.04 | | 7.51 | | 0.57 | | 3.14E-06 | | 4.80E-04 | |
| | | LRT | 98.66 | *** | 190.84 | *** | 234.17 | *** | 212.75 | *** | 201.24 | *** | 133.50 | *** | 145.22 | *** |
| | Block/Rep | Variance Component | 0.33 | | 5.55E-03 | | 0.00 | | 0.60 | | 0.05 | | 3.42E-07 | | 5.13E-05 | |
| | | LRT | 14.47 | *** | 8.36 | ** | 17.46 | *** | 11.08 | *** | 11.50 | *** | 14.43 | *** | 14.29 | *** |
| | Residual | Variance Component | 2.04 | | 0.03 | | 0.01 | | 2.51 | | 0.21 | | 1.74E-06 | | 2.40E-04 | |
| | | h$^2$ | 0.77 | | 0.85 | | 0.89 | | 0.87 | | 0.86 | | 0.81 | | 0.83 | |
| | Mean | | 19.00 | | 7.77 | | 6.87 | | 41.95 | | 23.02 | | 0.99 | | 1.13 | |
| | Accuracy | | 0.84 | | 0.92 | | 0.93 | | 0.93 | | 0.92 | | 0.89 | | 0.89 | |
| Joint Analysis | Inbred lines/G | Variance Component | 3.08 | | 0.10 | | 0.05 | | 8.40 | | 0.64 | | 3.70E-06 | | 5.65E-04 | |
| | | LRT | 176.37 | *** | 349.34 | *** | 369.33 | *** | 360.58 | *** | 355.46 | *** | 272.85 | *** | 289.54 | *** |
| | Block/Rep | Variance Component | 0.10 | | 1.80E-03 | | 2.92E-04 | | 0.10 | | 0.01 | | 1.56E-07 | | 2.20E-05 | |
| | | LRT | 7.05 | ** | 12.46 | *** | 2.54 | . | 6.98 | ** | 7.65 | ** | 29.85 | *** | 29.42 | *** |
| | Inbred lines/G X E | Variance Component | 0.13 | | 7.78E-05 | | 0.00 | | 0.06 | | 0.00 | | 4.40E-08 | | 3.46E-06 | |
| | | LRT | 1.16 | | 0.00 | | 0.04 | | 0.25 | | 0.04 | | 0.27 | | 0.08 | |
| | Residual | Variance Component | 2.13 | | 0.03 | | 0.01 | | 2.16 | | 0.17 | | 1.46E-06 | | 2.07E-04 | |
| | | h$^2$ | 0.74 | | 0.93 | | 0.94 | | 0.93 | | 0.93 | | 0.91 | | 0.91 | |
| | Mean | | 19.37 | | 7.80 | | 6.90 | | 42.31 | | 23.12 | | 0.99 | | 1.13 | |
| | Accuracy | | 0.86 | | 0.94 | | 0.94 | | 0.94 | | 0.94 | | 0.91 | | 0.92 | |

HSW: hundred-seed weight (g); L: seed length (mm); W: seed width (mm); AS: seed area (mm2); PS: seed perimeter (mm); CS: seed circularity; LWR: length/width ratio; Env.1: with fungicides for Asian soybean rust; Env. 2: without fungicides for Asian soybean rust; VC: Variance Component; LRT: Likelihood Ratio Test; H2: broad-sense heritability. '***','**','*', '.' Significant at the 0.0001, 0.001, 0.01, 0.05 probability levels, respectively. ns: not significant.

Supplementary Table 2. Analysis of deviance (ANADEV) for the hundred-seed weight and morphological characteristics of seeds, in each environment and in the joint analysis of the two environments. Samples of 100 seeds were obtained in experiments containing 16 parental lines and 3 checks designed in randomized blocks.

| Env. | Effects Random | | HSW | | L | | W | | AS | | PS | | CS | | LWR | |
|---|---|---|---|---|---|---|---|---|---|---|---|---|---|---|---|---|
| E1 | Parental lines/G | Variance Component | 4.03 | | 0.18 | | 0.08 | | 14.59 | | 1.15 | | 4.04E-06 | | 0.00 | |
| | | LRT | 13.07 | *** | 27.83 | *** | 29.50 | *** | 25.34 | *** | 25.79 | *** | 29.075 | *** | 31.68 | *** |
| | Block | Variance Component | 0.00 | | 1.17E-03 | | 1.05E-03 | | 0.18 | | 0.01 | | 5.92E-08 | | 0.00 | |
| | | LRT | 0.00 | | -1.17 | | -1.58 | | -5.37 | | -5.16 | | 16.243 | *** | 14.87 | *** |
| | Residual | Variance Component | 1.66 | | 0.02 | | 0.01 | | 2.26 | | 0.17 | | 4.85E-07 | | 0.00 | |
| | $h^2$ | | 0.83 | | 0.94 | | 0.94 | | 0.93 | | 0.93 | | 0.94 | | 0.95 | |
| | Mean | | 19.78 | | 7.78 | | 6.93 | | 42.33 | | 23.11 | | 1.00 | | 1.12 | |
| | Accuracy | | 0.91 | | 0.97 | | 0.97 | | 0.96 | | 0.96 | | 0.97 | | 0.98 | |
| E2 | Parental lines/G | Variance Component | 5.27 | | 0.17 | | 0.08 | | 13.33 | | 1.10 | | 3.77E-06 | | 7.01E-04 | |
| | | LRT | 11.40 | *** | 29.39 | *** | 31.08 | *** | 30.71 | *** | 30.95 | *** | 16.43 | *** | 20.759 | *** |
| | Block | Variance Component | 0.00 | | 1.50E-03 | | 0.00 | | 1.98E-07 | | 1.47E-08 | | 3.14E-07 | | 5.09E-05 | |
| | | LRT | 0.00 | | 0.39 | | 0.00 | | -7.97E-07 | | -1.54E-07 | | 3.59 | * | 3.9517 | * |
| | Residual | Variance Component | 2.54 | | 0.02 | | 0.01 | | 1.96 | | 0.15 | | 9.15E-07 | | 1.36E-04 | |
| | $h^2$ | | 0.81 | | 0.94 | | 0.94 | | 0.93 | | 0.94 | | 0.89 | | 0.91 | |
| | Mean | | 18.60 | | 7.67 | | 6.83 | | 41.20 | | 22.80 | | 0.99 | | 1.121438 | |
| | Accuracy | | 0.90 | | 0.97 | | 0.97 | | 0.97 | | 0.97 | | 0.94 | | 0.95 | |
| Joint Analysis | Parental lines/G | Variance Component | 3.57 | | 0.10 | | 0.05 | | 8.78 | | 0.67 | | 3.89E-06 | | 5.95E-04 | |
| | | LRT | 1.06E+01 | *** | 359.31 | *** | 402.63 | *** | 382.25 | *** | 376.86 | *** | 266.54 | *** | 282.21 | *** |
| | Block | Variance Component | 2.12E-07 | | 2.55E-03 | | 5.26E-04 | | 0.17 | | 0.01 | | 5.53E-08 | | 8.47E-06 | |
| | | LRT | -2.92E-06 | | 40.50 | *** | 19.18 | *** | 33.13 | *** | 33.86 | *** | 15.01 | *** | 16.54 | *** |
| | Parental lines/G X E | Variance Component | 0.62 | | 1.55E-03 | | 1.17E-09 | | 0.08 | | 4.94E-03 | | 1.85E-07 | | 2.68E-05 | |
| | | LRT | 1.16 | | 1.84 | . | -3.21E-06 | | 0.65 | | 0.46 | | 4.90 | * | 6.35 | ** |
| | Residual | Variance Component | 2.10 | | 0.03 | | 0.01 | | 2.21 | | 0.18 | | 1.47E-06 | | 2.08E-04 | |
| | $h^2$ | | 0.79 | | 0.93 | | 0.94 | | 0.94 | | 0.94 | | 0.91 | | 0.92 | |
| | Mean | | 19.14 | | 7.80 | | 6.90 | | 42.30 | | 23.12 | | 0.99 | | 1.13 | |
| | Accuracy | | 0.88 | | 0.94 | | 0.94 | | 0.94 | | 0.94 | | 0.92 | | 0.92 | |

HSW: hundred-seed weight (g); L: seed length (mm); W: seed width (mm); AS: seed area (mm2); PS: seed perimeter (mm); CS: seed circularity; LWR: length/width ratio; Env.1: with fungicides for Asian soybean rust; Env. 2: without fungicides for Asian soybean rust; VC: Variance Component; LRT: Likelihood Ratio Test; H2: broad-sense heritability. '***','**','*', '.' Significant at the 0.0001, 0.001, 0.01, 0.05 probability levels, respectively. ns: not significant.

**SUPPLEMENTARY FIGURES**

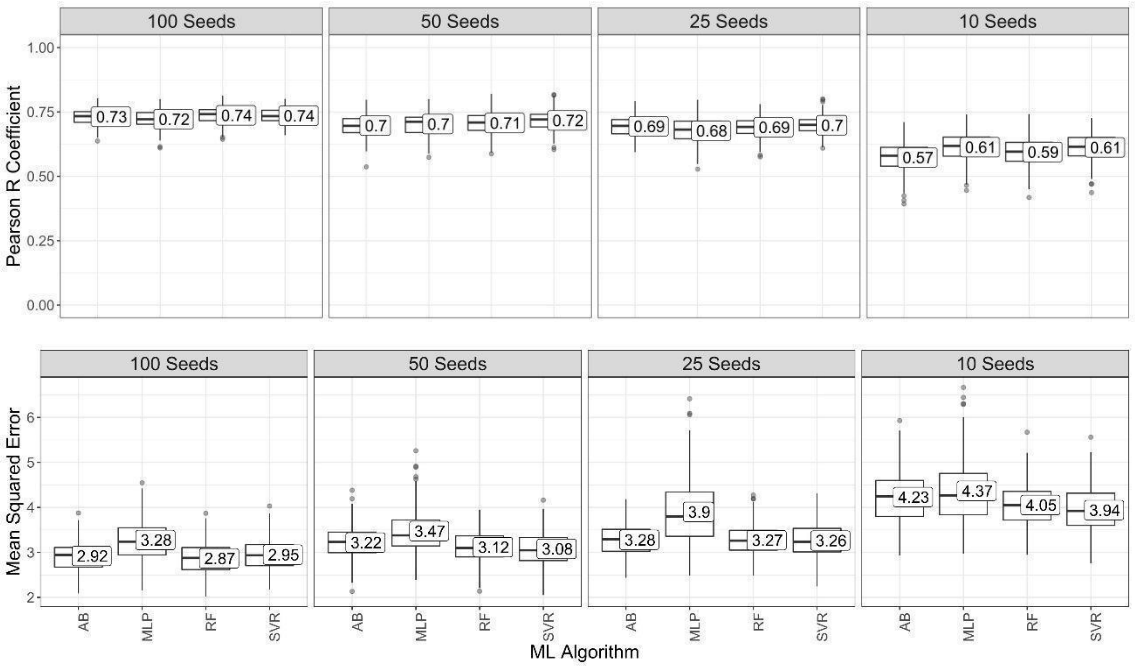


Supplementary Fig. 1. Predictive performance evaluation of machine learning (ML) algorithms (*AdaBoost*, *multilayer perceptron* (*MLP*), *random forest* (*RF*), and *support vector regression* (*SVR*)) for prediction of the hundred-seed weights using the images of the environment 1 considering the Pearson R coefficient and the mean squared error as evaluation metrics.

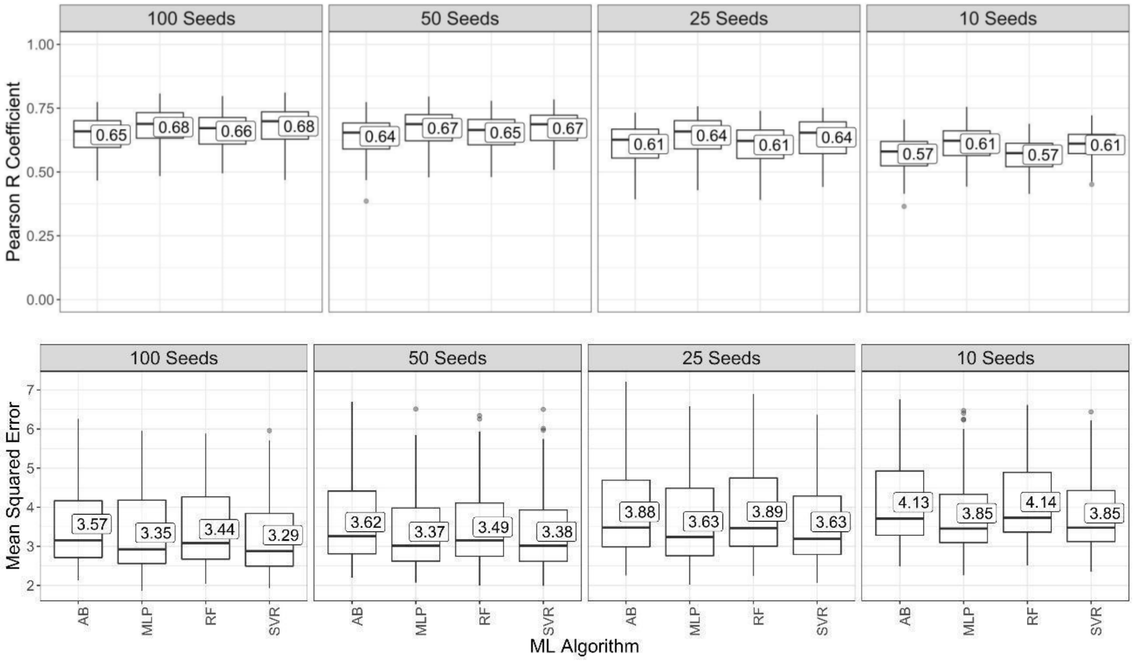


Supplementary Fig. 2. Predictive performance evaluation of machine learning (ML) algorithms (*AdaBoost*, *multilayer perceptron* (*MLP*), *random forest* (*RF*), and *support vector regression* (*SVR*)) for prediction of the hundred-seed weights using the images of the environment 2 considering the Pearson R coefficient and the mean squared error as evaluation metrics.
