## Supplementary File 1 for "A novel image-based approach for soybean seed phenotyping using machine learning techniques"

### SegmentationCode_Example

October 10, 2022

[9]:
```
#Libraries
from skimage import io as skio
from skimage import filters
import matplotlib.pyplot as plt
import cv2
import os
import numpy as np
import pandas as pd
```

[2]:
```
#Reading image
img1 = cv2.imread("42.371-426D.jpg", 0)
img3 = cv2.imread("42.371-426D.jpg", 1)

fig = plt.figure(figsize=(20,10))
plt.imshow(img3)
```

[2]:
```
<matplotlib.image.AxesImage at 0x7f0a54b21910>
```

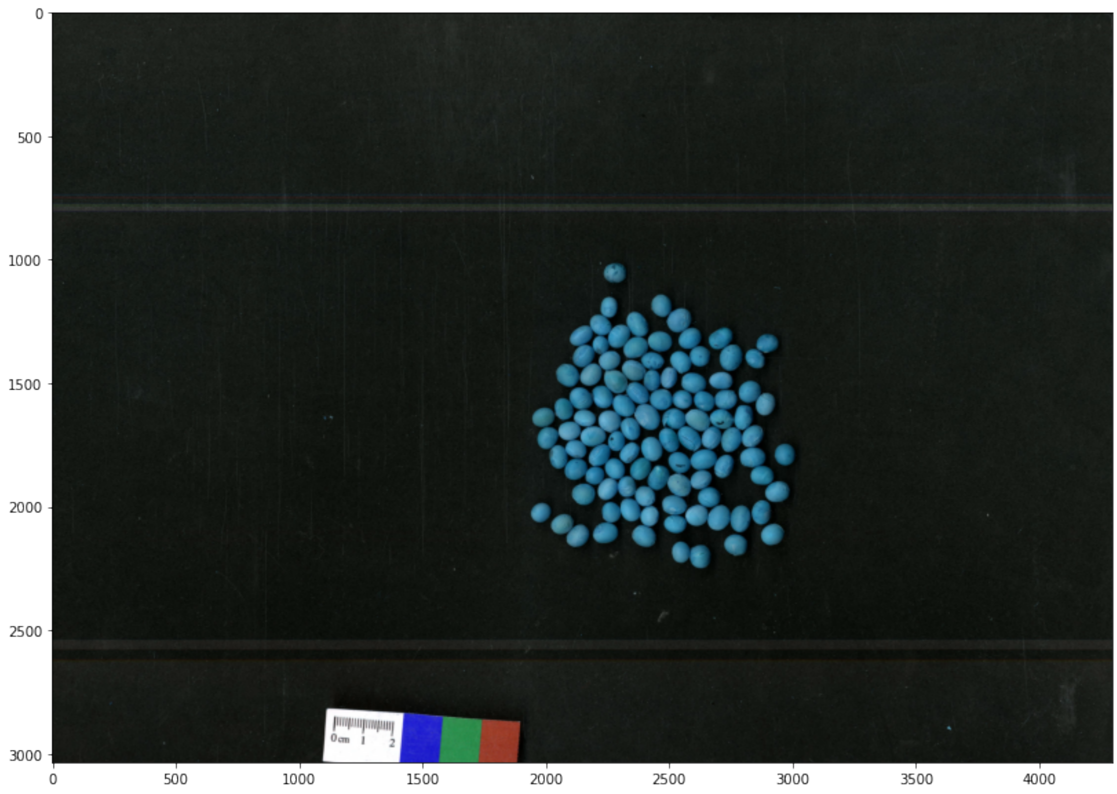


[3]:
```
#Erode the components
kernel1 = np.ones((25,25),np.uint8)
erosion = cv2.erode(img3,kernel1,iterations=1)

fig = plt.figure(figsize=(20,10))
plt.imshow(erosion)
```

[3]: <matplotlib.image.AxesImage at 0x7f0a54a3c9d0>

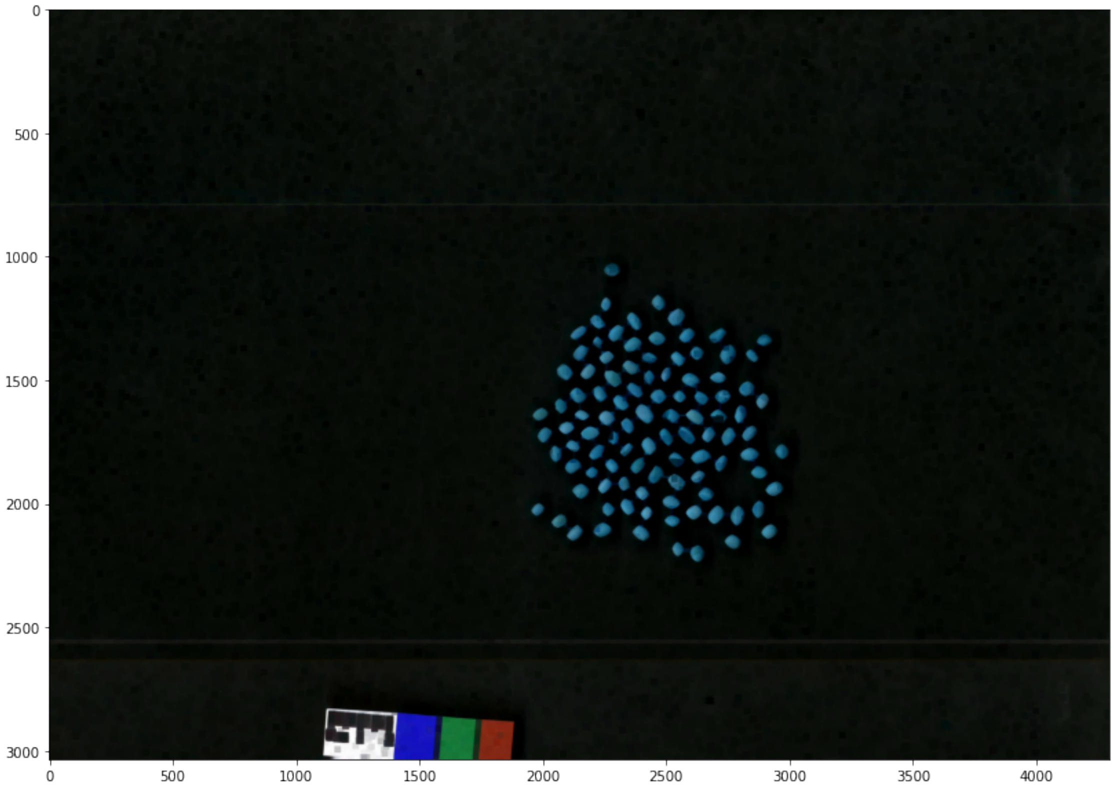


[4]:
```
#Closing the components
kernel2 = np.ones((11,11),np.uint8)
closing = cv2.morphologyEx(erosion,cv2.MORPH_CLOSE, kernel2)

fig = plt.figure(figsize=(20,10))
plt.imshow(closing)
```

[4]: <matplotlib.image.AxesImage at 0x7f0a541b3970>

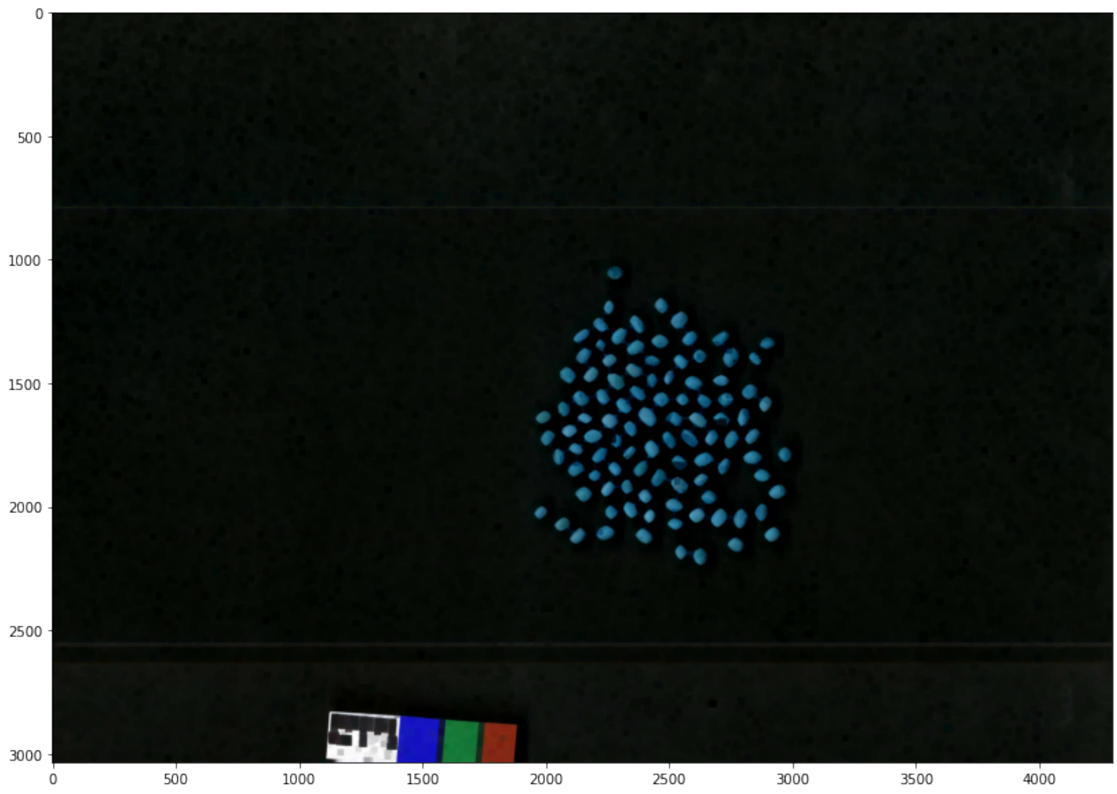

```
[5]: #Lighting image
     alpha = 2.5
     beta = 50

     adjusted = cv2.convertScaleAbs(closing,alpha=alpha,beta=beta)
     adjusted1 = cv2.cvtColor(adjusted, cv2.COLOR_BGR2GRAY)

     fig = plt.figure(figsize=(20,10))
     plt.imshow(adjusted1, cmap="gray")
```

```
[5]: <matplotlib.image.AxesImage at 0x7f0a4c1e58b0>
```

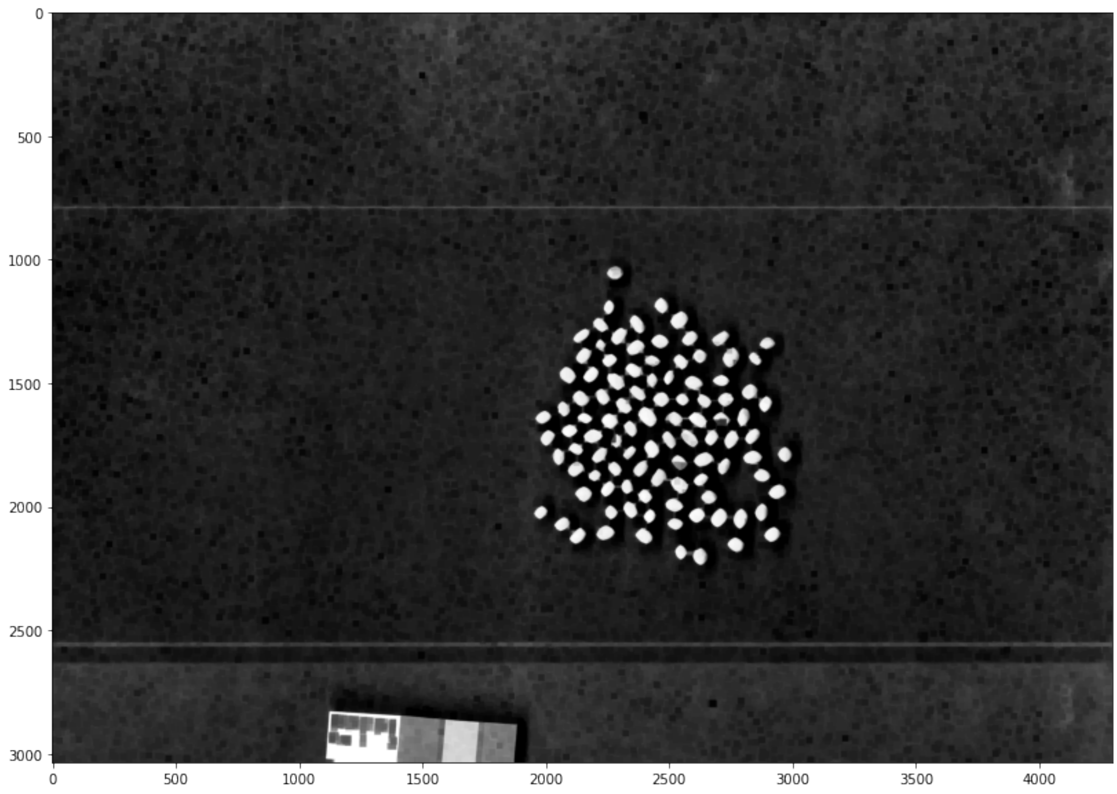

[6]:
```
#Find components
blur2 = cv2.blur(adjusted1,(25,25),0)
ret, th = cv2.threshold(blur2,0,255, cv2.THRESH_BINARY+cv2.THRESH_OTSU)
num_labels, labels = cv2.connectedComponents(th)

fig = plt.figure(figsize=(20,10))
plt.imshow(th, cmap="gray")
```

[6]: <matplotlib.image.AxesImage at 0x7f0a4c14d4c0>

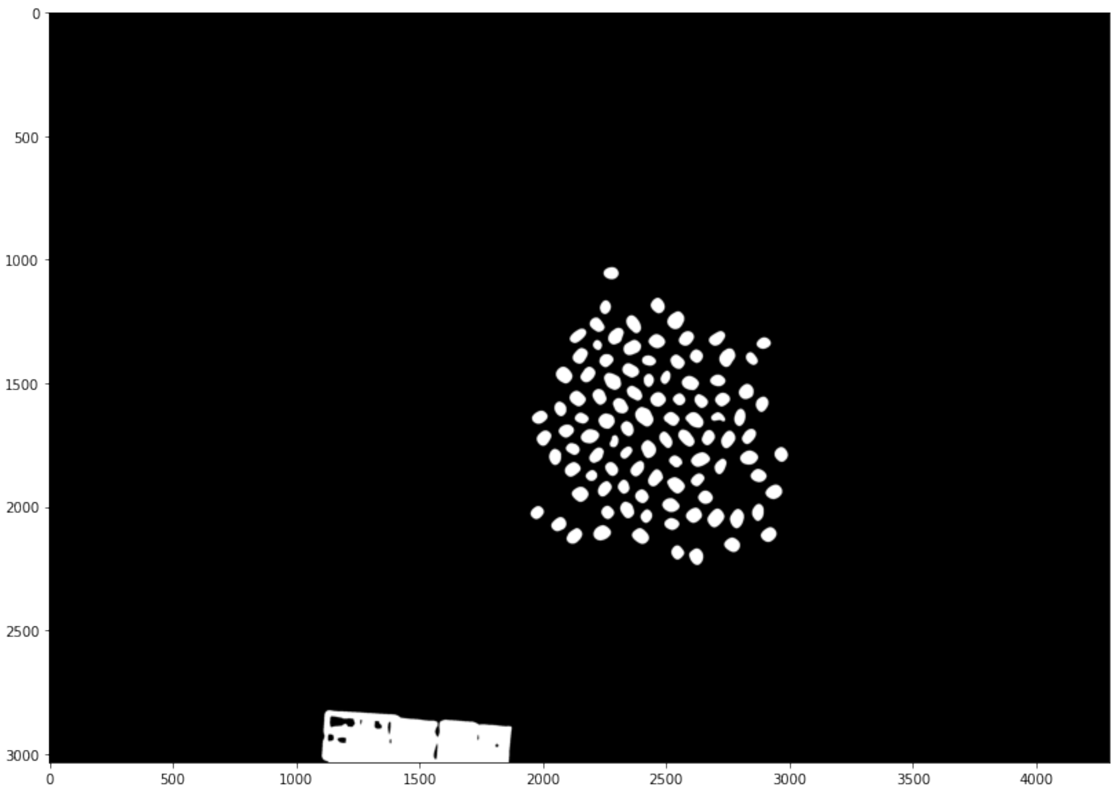


[7]:
```
#Selecting only the seeds
j = 0
qtd = 0

#Going component by component
for i in range(1,num_labels):
    flag = 0

    #How many pixels?
    aux_qtd = np.count_nonzero(labels == i)

    #Check a range size
    if((aux_qtd < 500)|(aux_qtd>8000)):
        labels[labels==i] = 0
        flag = 1

    #Found a seed!
    else:
        qtd += 1

    #Updating labels
    labels[labels==i] = labels[labels==i]-j
```

```
    if(flag):
        j += 1

th[labels == 0] = 0

fig = plt.figure(figsize=(20,10))
plt.imshow(th, cmap="gray")
```

[7]: <matplotlib.image.AxesImage at 0x7f0a4c13b9d0>

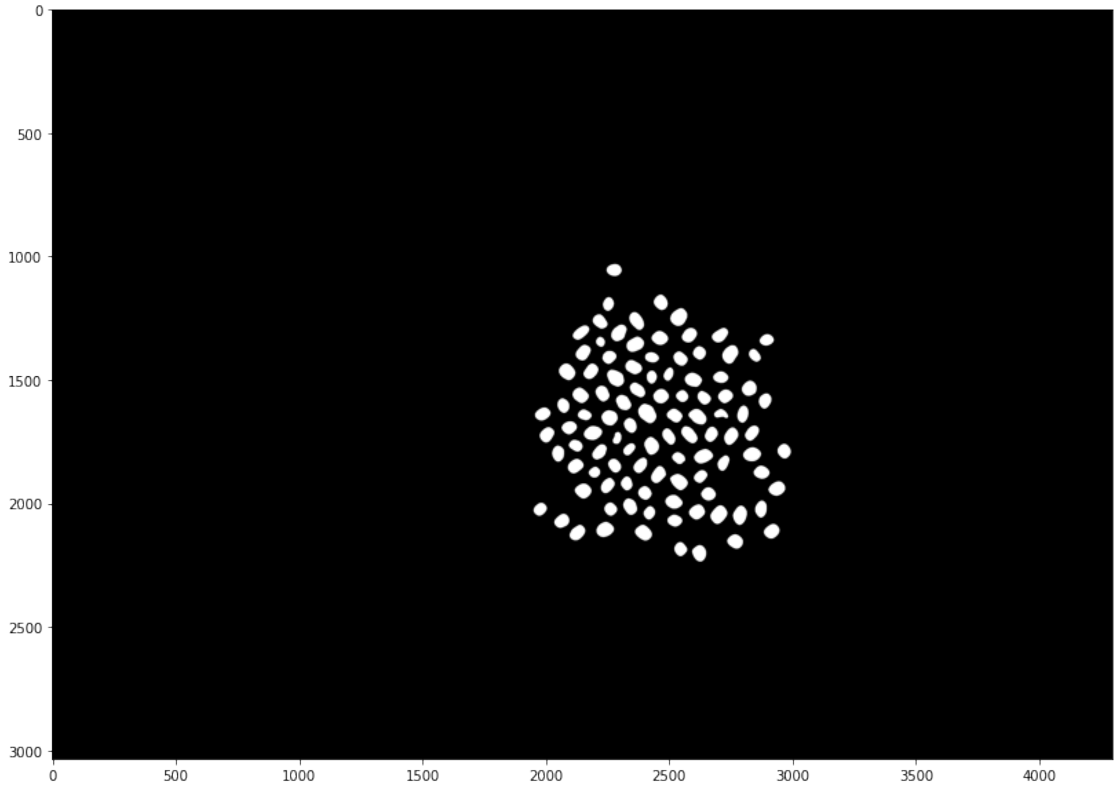


[8]:
```
#Taking the contour of segmentation
cont = cv2.findContours(th,cv2.RETR_EXTERNAL,
                    cv2.CHAIN_APPROX_SIMPLE)[0]

#Showing contour
img_contour = img3.copy()
img_contour = cv2.drawContours(img_contour,cont,-1,(255,0,0),15)

fig = plt.figure(figsize=(20,10))
plt.imshow(img_contour)
```

[8]: <matplotlib.image.AxesImage at 0x7f0a4c0ae580>

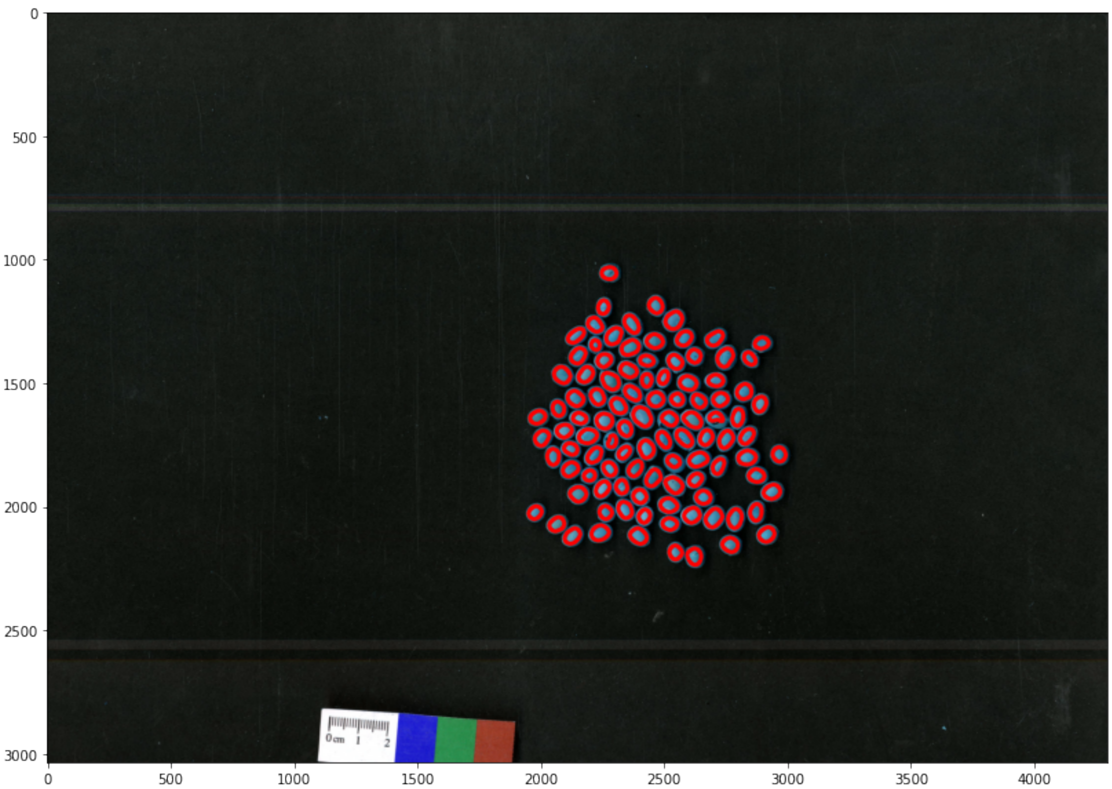
0
500
1000
1500
2000
2500
3000
0
500
1000
1500
2000
2500
3000
3500
4000
0 cm 1 2
