## Supplementary File 2 for "A novel image-based approach for soybean seed phenotyping using machine learning techniques"

### SegmentationCode

October 10, 2022

[7]:

```
#Libraries
from skimage import io as skio
from skimage import filters
import matplotlib.pyplot as plt
import cv2
import os
import numpy as np
import pandas as pd
```

[2]:

```
#Identifying components
def components(img,img_3,output):
    #Erode the components
    kernel1 = np.ones((25,25),np.uint8)
    erosion = cv2.erode(img_3,kernel1,iterations=1)

    #Closing the components
    kernel2 = np.ones((11,11),np.uint8)
    closing = cv2.morphologyEx(erosion,cv2.MORPH_CLOSE, kernel2)

    #Lighting image
    alpha = 2.5
    beta = 50

    adjusted = cv2.convertScaleAbs(closing,alpha=alpha,beta=beta)
    adjusted1 = cv2.cvtColor(adjusted, cv2.COLOR_BGR2GRAY)

    #Find components
    blur2 = cv2.blur(adjusted1,(25,25),0)
    ret, th = cv2.threshold(blur2,0,255, cv2.THRESH_BINARY+cv2.THRESH_OTSU)
    num_labels, labels = cv2.connectedComponents(th)

    #Selecting only the seeds
    j = 0
    qtd = 0

    #Going component by component
    for i in range(1,num_labels):
```

```
        flag = 0

        #How many pixels?
        aux_qtd = np.count_nonzero(labels == i)

        #Check a range size
        if((aux_qtd < 500)|(aux_qtd>8000)):
            labels[labels==i] = 0
            flag = 1

        #Found a seed!
        else:
            qtd += 1

        #Updating labels
        labels[labels==i] = labels[labels==i]-j
        if(flag):
            j += 1

    th[labels == 0] = 0

    #Ellipse sizes
    sizes = list()

    #Taking the image cuts
    for i in range(1,qtd+1):

        #Making a copy
        img_aux = img_3.copy()
        th2 = th.copy()
        th2[labels != i] = 0

        #Dilating the components
        kernel = np.ones((25,25),np.uint8)
        th_dil = cv2.dilate(th2,kernel,iterations=1)

        #Selecting the seed region
        img_aux[th_dil == 0] = [0,0,0]

        #Taking the contour of segmentation
        cont = cv2.findContours(th_dil,cv2.RETR_EXTERNAL,
                                cv2.CHAIN_APPROX_SIMPLE)[0]

        #Finding the ellipse
        ellipse = cv2.fitEllipse(cont[0])
        sizes.append(ellipse)
        img_aux2 = img_aux.copy()
```

```
        img_aux_ellipse = img_aux.copy()

        #Showing ellipse
        img_aux2 = cv2.ellipse(img_aux2, ellipse, (0, 0, 255),3)
        img_aux_ellipse = cv2.ellipse(img_aux_ellipse, ellipse, (255, 255,␣
↪255),thickness=-1)

        #Cutting ellipse
        img_ellipse = img_3.copy()
        img_ellipse = np.bitwise_and(img_ellipse,img_aux_ellipse)

        #Showing contour
        img_contour = img_3.copy()
        img_contour = cv2.drawContours(img_contour,cont,-1,(255,0,0),5)

        #Defining minimum area around it
        rect = cv2.minAreaRect(cont[0])
        box = cv2.boxPoints(rect)
        box = np.int0(box)

        #Expanding 10 pixels
        box[0] = box[0] - 10
        box[1,0] = box[1,0] + 10
        box[1,1] = box[1,1] - 10
        box[2] = box[2] + 10
        box[3,0] = box[3,0] - 10
        box[3,1] = box[3,1] + 10

        #Cropping the image
        width = int(rect[1][0])
        height = int(rect[1][1])

        #Points
        src_pts = box.astype("float32")
        dst_pts = np.array([[0,height-1],[0,0],
                        [width-1,0],[width-1,height-1]],
                        dtype="float32")

        #Cutting
        M = cv2.getPerspectiveTransform(src_pts,dst_pts)
        warped0 = cv2.warpPerspective(img_3,M,(width,height))
        warped1 = cv2.warpPerspective(img_aux,M,(width,height))
        warped2 = cv2.warpPerspective(img_contour,M,(width,height))
        warped3 = cv2.warpPerspective(img_aux2,M,(width,height))
        warped4 = cv2.warpPerspective(img_ellipse,M,(width,height))
        cv2.imwrite(output+"/"+str(i)+"_0.jpg",warped0)
        cv2.imwrite(output+"/"+str(i)+"_1.jpg",warped1)
```

```
        cv2.imwrite(output+"/"+str(i)+"_2.jpg",warped2)
        cv2.imwrite(output+"/"+str(i)+"_3.jpg",warped3)
        cv2.imwrite(output+"/"+str(i)+"_4.jpg",warped4)

    return(sizes)
```

[3]:

```
#Calculate measures
def calculate(ellipse,name):
    length_aux = max(ellipse[1])/2 #Comprimento
    width_aux = min(ellipse[1])/2 #Largura

    length_aux = length_aux * 0.08474576
    width_aux = width_aux * 0.08474576

    #Area
    area = length_aux * width_aux * np.pi

    #Perimeter
    a = length_aux
    b = width_aux
    perimeter = np.pi*((a+b)+((3*(a-b)**2)/(10*(a+b)+np.
 ↪sqrt(a**2+14*a*b+b**2))))

    #Ratio length/width
    ratio = length_aux/width_aux

    #Circularity
    circ = 4*np.pi * area/(perimeter**2)

    measures = pd.DataFrame({
                    "Area":[area],
                    "Perimeter":[perimeter],
                    "Ratio":[ratio],
                    "Circularity":[circ],
                    "Length":[length_aux*2],
                    "Width":[width_aux*2],
                    "MinorAxis":[width_aux],
                    "Major":[length_aux],
                    "Photo":[name]})

    return(measures)
```

[4]:

```
#Reading directory
files = os.listdir(".")
final_table = pd.DataFrame()

for i in files:
```

```
    if "jpg" in i:
        img1 = cv2.imread(i, 0)
        img3 = cv2.imread(i, 1)

        #Create an image output directory
        os.makedirs("Segmentation/" + i)

        #Taking measures
        calculations = components(img1,img3,"Segmentation/"+i)

        for j in calculations:
            final_table = pd.concat([final_table,calculate(j,i)])
```

[5]:
```
final_table
```

[5]:
```
         Area  Perimeter     Ratio  Circularity    Length     Width  \
0   41.974661  23.100235  1.192537     0.988472  7.983340  6.694416
0   44.550021  23.707411  1.107971     0.996069  7.927627  7.155085
0   36.461157  21.549883  1.209033     0.986621  7.491854  6.196569
0   55.687832  26.532110  1.134046     0.994092  8.967068  7.907149
0   49.770516  25.444106  1.356146     0.966068  9.270313  6.835776
..        …          …         …          …         …         …
0   47.162795  24.553409  1.238370     0.983073  8.623429  6.963533
0   50.338629  25.256963  1.161689     0.991628  8.628807  7.427809
0   46.499898  24.228989  1.117528     0.995385  8.134116  7.278666
0   39.751707  22.386672  1.097676     0.996751  7.453669  6.790407
0   47.664894  24.556331  1.143360     0.993303  8.330018  7.285558

    MinorAxis     Major            Photo
0    3.347208  3.991670  42.371-426D.jpg
0    3.577543  3.963814  42.371-426D.jpg
0    3.098284  3.745927  42.371-426D.jpg
0    3.953575  4.483534  42.371-426D.jpg
0    3.417888  4.635157  42.371-426D.jpg
..        …         …              …
0    3.481767  4.311715  42.371-426D.jpg
0    3.713904  4.314404  42.371-426D.jpg
0    3.639333  4.067058  42.371-426D.jpg
0    3.395203  3.726835  42.371-426D.jpg
0    3.642779  4.165009  42.371-426D.jpg

[100 rows x 9 columns]
```

[6]:
```
final_table.to_csv("Measures.csv")
```
